# On the Origin of Obligate Parthenogenesis in *Daphnia pulex*

**DOI:** 10.1101/2022.10.10.511613

**Authors:** Marelize Snyman, Sen Xu

**Affiliations:** Department of Biology, University of Texas at Arlington, Arlington, Texas, 76019, USA

**Keywords:** *Daphnia*, obligate parthenogenesis, meiosis, cell cycle

## Abstract

Despite the presence of obligately parthenogenetic (OP) lineages derived from sexual ancestors in diverse phylogenetic groups, the genetic mechanisms giving rise to the OP lineages remain poorly understood. The freshwater microcrustacean *Daphnia pulex* typically reproduces via cyclical parthenogenesis. However, some populations of OP *D. pulex* have emerged due to ancestral hybridization and introgression events between two cyclically parthenogenetic (CP) species *D. pulex* and *D. pulicaria*. These OP hybrids produce both subitaneous and resting eggs parthenogenetically, deviating from CP isolates where resting eggs are produced via conventional meiosis and mating. This study examines the genome-wide expression and alternative splicing patterns of early subitaneous versus early resting egg production in OP *D. pulex* isolates to gain insight into the genes and mechanisms underlying this transition to obligate parthenogenesis. Our differential expression and functional enrichment analyses revealed a downregulation of meiosis and cell cycle genes during early resting egg production, as well as divergent expression patterns of metabolism, biosynthesis, and signaling pathways between the two reproductive modes. These results provide important gene candidates for future experimental verification, including the CDC20 gene that activates the anaphase-promoting complex in meiosis.

## Introduction

The existence of obligately parthenogenetic eukaryotic lineages that have entirely abandoned sexual reproduction has long fascinated evolutionary biologists. The transition from sexual reproduction to obligate parthenogenesis is phylogenetically widespread, having occurred independently in most multicellular taxa (Bell 2019, Neiman *et al*. 2014). Various cytogenetic manifestations of obligate parthenogenesis (i.e., modifications of meiosis) have been well described in the literature, including automictic parthenogenesis, apomictic parthenogenesis, gynogenesis, and hybridogenesis (Stenberg and Saura 2009, Vrijenhoek 1998). However, the molecular mechanisms and genes underlying these cytogenetic modifications are poorly understood (Ferree *et al*. 2006, King and Hurst 2010, Riparbelli *et al*. 2005, Suomalainen *et al*. 1987).

Obligate parthenogenesis can originate through multiple evolutionary routes. Spontaneous mutations in genes involved in meiosis or reproduction could lead to the loss of sexual reproduction as in monogonont rotifers (Serra and Snell 2009). Contagious parthenogenesis could result due to the spread of asexuality conferring-elements as in the pea aphid (Jaquiéry *et al*. 2014), and parasite-induced parthenogenesis could occur for example, in wasps infected by *Wolbachia* (Simon *et al*. 2003). The most common route, interspecific hybridization, could disrupt meiosis due to genetic incompatibilities between parental species, resulting in the loss of sex (Vrijenhoek 1998, White 1978). Among vertebrates, there are currently around 100 known parthenogenetic lineages of amphibians, reptiles, and fish that are interspecific hybrids (Avise 2008, Avise 2015, Dawley and Bogart 1989, Neaves and Baumann 2011). For invertebrates, the occurrence of hybrid parthenogens have been demonstrated in snails (Johnson and Bragg 1999), crustaceans (Innes and Hebert 1988), and many insects (Schwander *et al*. 2011, Stenberg and Lundmark 2004, White *et al*. 1977).

This widespread occurrence of obligate parthenogens with a hybrid ancestry suggests that understanding meiotic modifications associated with hybridization may provide insight into the origin of obligately parthenogenetic (OP) species. In this study, we investigate the origin of obligate parthenogenesis in the cladoceran microcrustacean, *Daphnia pulex*, commonly found in freshwater habitats in North America. Previous studies have shown that obligately parthenogenetic *Daphnia pulex* originated due to hybridization and introgression events between two cyclically parthenogenetic (CP) sister species, *D. pulex* and *D. pulicaria* (Xu *et al*. 2013, 2015). This hypothesis is supported by genome-wide association studies showing microsatellite and SNP alleles exclusively associated with OP *D*.*pulex* originated in *D. pulicaria* (Tucker *et al*. 2013, Xu *et al*. 2013, 2015). To date, no reproductive mode tests performed on *D. pulicaria* isolates have revealed OP lineages (Heier and Dudycha 2009). Having diverged about 800,000 – 2,000,000 years ago (Colbourne and Hebert 1996, Cristescu *et al*. 2012, Omilian and Lynch 2009), CP *D. pulex* and *D. pulicaria*, members of the *D. pulex* species complex, are morphologically similar (Brandlova *et al*. 1972), yet can be distinguished due to inhabiting very different environments and by species-specific microsatellite markers or allozyme loci such as the lactate dehydrogenase (Ldh) locus (Cristescu *et al*. 2014).

The main difference between the cyclically parthenogenetic parental species and obligately parthenogenetic hybrids is how they reproduce. Under favorable environmental conditions such as high food abundance and low population density, both CP and OP *Daphnia* females (**Figure 1A, Figure 1B**) reproduce asexually, producing diploid subitaneous eggs through a modified meiosis. In this modified meiosis, during meiosis I, bivalents align at the metaphase plate; however, cell division is arrested before the onset of anaphase I, after which each half-bivalent moves back to the metaphase plate and sister chromatids rearrange (Hiruta *et al*. 2010). Thus, during modified meiosis I, there is no segregation of homologous chromosomes and no cytokinesis resulting in daughter cells. Next, meiosis II proceeds normally and results in diploid embryos (Ojima 1958, Zaffagnini and Sabelli 1972, Hiruta *et al*. 2010). In the absence of rare events such as ameiotic recombination, conversion, and mutation (Omilian *et al*. 2006, Xu *et al*. 2009), all asexually produced progenies, mostly female, are genetically identical to the mother.

**Figure 1.**
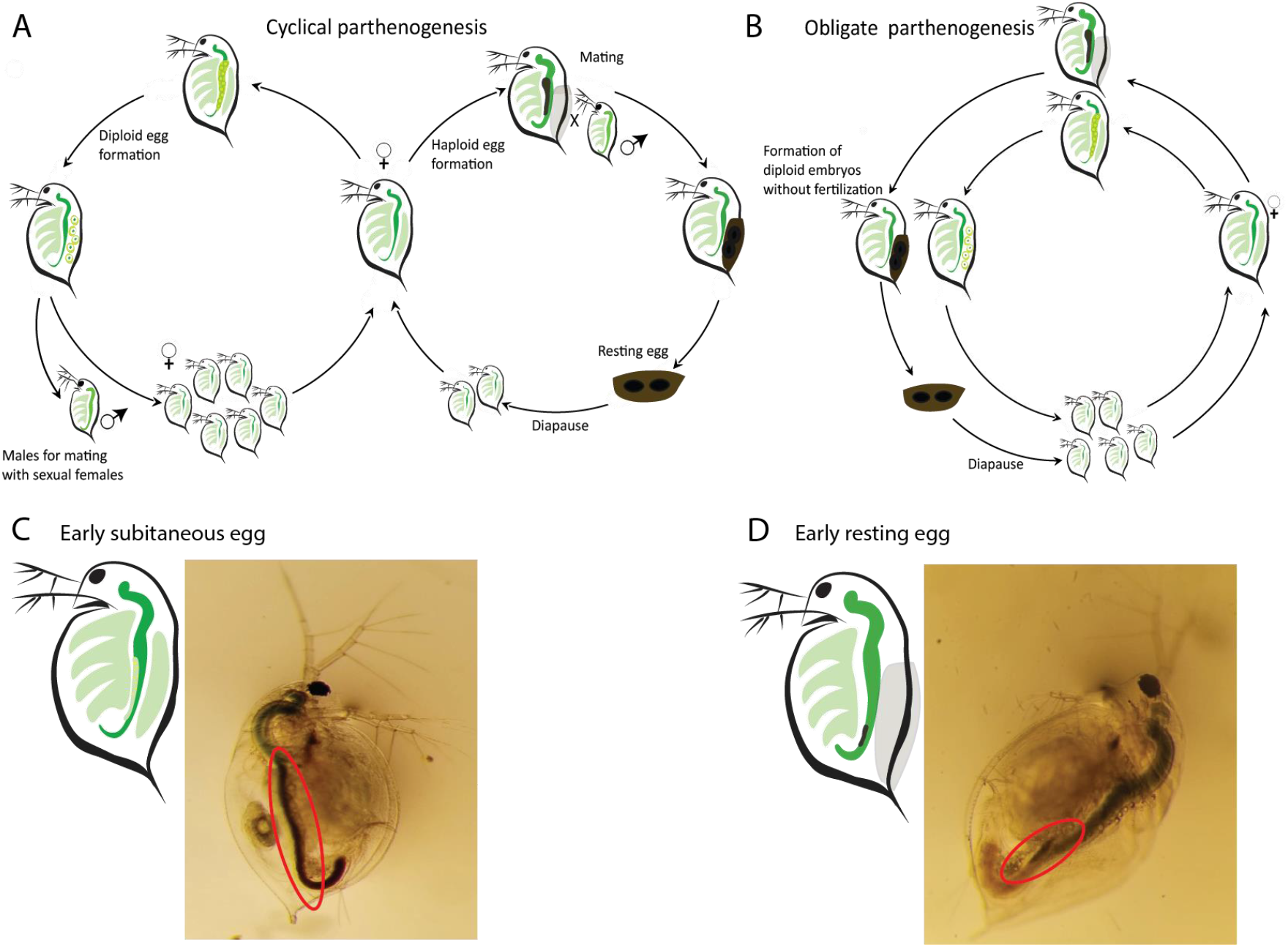
(A) Cyclically parthenogenetic and (B) obligately parthenogenetic life cycles in *Daphnia*. (C) Early subitaneous egg and (D) early resting egg production as determined by the color and size of the ovaries (red circles).

Once the environment deteriorates (e.g., low food abundance, high population density), some subitaneous embryos can develop into males through environmental sex determination (Gorr *et al*. 2006, Tatarazako *et al*. 2003). The CP *Daphnia* females switch to produce haploid eggs (usually two) through meiosis, which upon fertilization by sperm become resting embryos. In contrast, OP *D. pulex* females produce chromosomally unreduced resting embryos without mating through a modified meiosis. Previous cytological observations have shown that parthenogenesis producing subitaneous eggs in OP and CP isolates is highly similar to parthenogenesis producing resting eggs in OP isolates (Zaffagnini and Sabelli 1972). The resulting resting embryos from OP isolates (without fertilization) are deposited into a protective case (i.e., ephippium) and remain dormant until environmental conditions turn favorable for hatching. As environmental conditions drive the alteration of reproductive modes in *Daphnia*, we argue that transcriptomic changes due to environmental variation may play a fundamental role and provide insight into the origin of obligate parthenogenesis.

Attempting to uncover the molecular basis of parthenogenesis, previous work by Xu *et al*. in 2015 functionally annotated the 647 asexual-specific SNPs introgressed from *D. pulicaria* into the *D. pulex* genome and identified 206 protein-coding genes associated with obligate parthenogenesis. Of these genes, 52 were predicted to have general or unknown functions and the analysis showed no functional enrichment for any pathway. The authors identified some candidate genes affecting meiosis, the cell cycle and DNA replication; however, they were unable to distinguish between causal genes or spurious associations (Xu *et al*. 2015). Building on this work, other studies have examined genome-wide expression differences between reproductive stages and modes to elucidate the role of gene expression changes in the origin of OP *Daphnia*. Xu *et al*. 2022 examined the genome-wide expression patterns during early resting egg production stage between various OP *D. pulex* and parental CP *D. pulex* and *D. pulicaria* isolates **(Figure 2A)**. The results showed that early resting egg production in OP *D. pulex* is associated with a downregulation of many meiosis and cell cycle genes, and an upregulation of metabolic and biosynthesis genes compared to early resting egg production in CP isolates (Xu *et al*. 2022). These results suggests that the origin of obligate parthenogenesis from meiosis is likely caused by the downregulation of key meiosis and cell cycle genes, and an upregulation of metabolism. Interestingly, when comparing the gene expression patterns between early subitaneous egg and early resting egg production within individual CP *D. pulex* and *D. pulicaria* isolates **(Figure 2B)**, meiosis and cell cycle genes were also found to be downregulated while metabolic, and biosynthesis genes were upregulated in early subitaneous egg development (i.e., parthenogenesis) compared to resting egg production (i.e., meiosis) in both species (Huynh *et al*. 2021). Together, the results from these two transcriptomic studies strongly suggest that parthenogenesis in both OP (resting egg production) and CP (subitaneous egg production) isolates are associated with a downregulation of meiosis and cell cycle genes and an upregulation of metabolic and biosynthesis genes, which may play essential roles in triggering parthenogenesis. Therefore, it has been hypothesized that the origin of obligate parthenogenesis in *Daphnia* involves the use of existing ameiotic germline cell division pathways (normally used in the production of asexual, subitaneous eggs) in the production of resting eggs (Xu *et al*. 2022).

**Figure 2.**
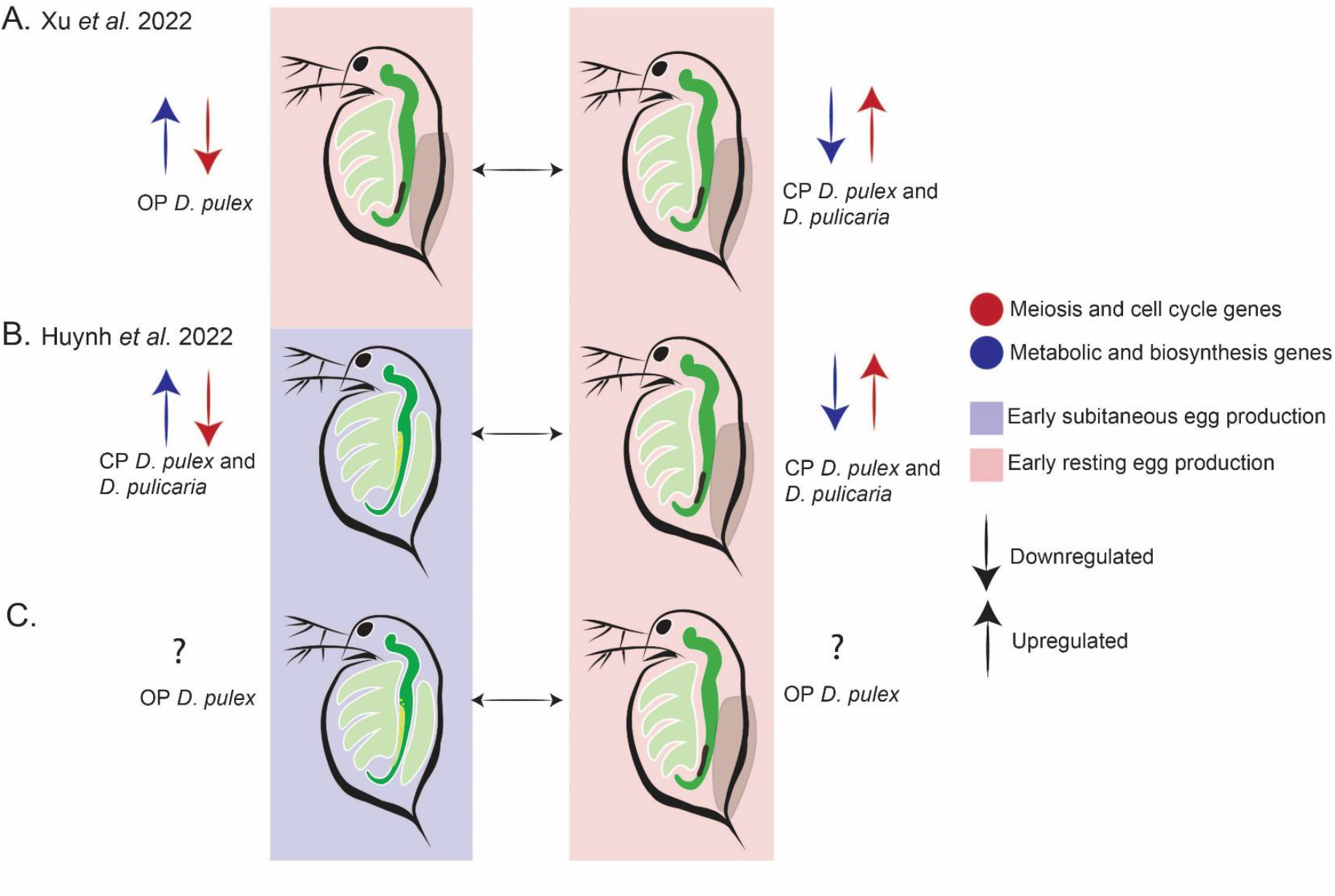
(A) Regulation of genes during early resting egg production between OP *D. pulex* and CP *D. pulex* and *D. pulicaria* isolates. (B) Regulation of genes during early resting egg production and early subitaneous egg production within CP isolates, (C) and within OP *D. pulex* isolates.

To further test this hypothesis, this work examines the genome-wide gene expression variation differentiating the early stages of two modes of parthenogenesis (subitaneous vs. resting eggs) within hybrid OP *D. pulex* isolates **(Figure 2C)**. We hope that these data will illuminate the genetic basis of the cytological modifications occurring in modified meiosis I, providing insight into the origin of obligate parthenogenesis. Furthermore, we set out to identify patterns of alternative splicing in the two reproductive modes to understand whether alternative splicing plays a role in the origin of obligate parthenogenesis.

## Materials and Methods

### Daphnia Isolates

A total of three obligate parthenogenetic (OP) *Daphnia pulex* isolates (DB4-4, K09, and Main 348-1) were used in this study. These isolates were previously collected from Texas, Ontario, and Maine, respectively **(Supplementary Table S1)** and have been maintained in the lab as clonal cultures. Initiated from a single female, each isolate has been kept as an asexually reproducing line in artificial lake water (Kilham *et al*. 1998) under a 16:8 light/dark cycle at 18°C and fed with the green algae, *Scenedesmus obliquus*, twice a week.

### Animal tissue collection

For each OP *D. pulex* isolate, experimental animals were maintained in the same environmental conditions for two generations to minimize maternal effects, which could significantly impact gene expression. Then, one-day-old neonates were continuously collected from each isolate and grown until sexual maturity in the same environmental conditions described above. Sexually mature animals were examined daily under a light microscope to collect females engaging in the early stage of producing subitaneous eggs and in the early stage of producing resting eggs. Early subitaneous embryo production is characterized by a thin clear, or greenish ovary extending along the gut with oil droplets visible. In contrast, early resting egg production is characterized by a small milky brown ovary starting to develop along the end of the gut **(Figure 1C and D)**. For each isolate, three replicates of each stage were collected (15-20 individuals) for RNA extraction.

### RNA extraction and sequencing

RNA of all samples was extracted using the Promega (Madison, WI, USA) SV Total RNA Isolation kit, following the manufacturer’s instructions. RNA concentration was measured using a Qubit 4.0 Fluorometer (Thermo Scientific, Waltham, MA, USA). RNA integrity was checked with a Bioanalyzer (Agilent Technologies, Santa Clara, CA, USA). RNA sequencing libraries were prepared by Novogene Corporation Inc. (Sacramento, CA, USA) using standard Illumina sequencing library protocol. Each library was sequenced on an Illumina NovaSeq6000 platform with at least 20 million 150-bp paired-end reads. The raw RNA sequence data were deposited at NCBI SRA under PRJNA847604.

### Sequencing quality control and mapping

The software package FastQC (Andrews 2010) was used to examine the quality of the raw reads. Because of no observed adapter contamination, reads were mapped directly to the *D. pulex* reference genome PA42 3.0 (Ye *et al*. 2017) using STAR aligner (Dobin *et al*. 2013) with default parameters. SAMtools (Li *et al*. 2009) was used to remove reads that mapped to multiple locations in the genome, and the program featureCounts (Liao *et al*. 2014) was used to obtain the raw counts for expressed genes in each sample.

### Differential gene expression analysis

We performed differential expression (DE) analysis using DESeq2 v.1.34.0 (Love *et al*. 2014) in R (R Core Team 2020). Differentially expressed genes (DEG) were determined for early subitaneous *vs* early resting egg development for each isolate individually to investigate for differential expression within each *Daphnia* isolate and by pooling all the isolates to establish commonalities. The Wald negative binomial test using the design formula ∼ Stage for individual isolates and ∼ Isolate + Stage for pooled isolates were utilized in DEseq2. P-values were adjusted for multiple testing using the Benjamini-Hochberg method, and genes were considered significantly differentially expressed if they had a p-value < 0.05 and a fold-change > 1.5 and < - 1.5 for upregulation and downregulation, respectively.

### KEGG (Kyoto Encyclopedia of Genes and Genomes) pathway and GO term enrichment analysis

To investigate the biological relevance of the differentially expressed genes, we performed a functional enrichment analysis using the R package topGO (Alexa and Rahnenfuhrer 2016). The default algorithm, weight01, was used along with the Fisher exact test, and GO terms were considered significantly enriched if the p-value was less than 0.05. Our script is available at https://github.com/Marelize007/RNAseq_obligate_parthenogenesis.

We also examined whether any KEGG pathways were enriched for differentially expressed genes. We queried 18,440 gene sequences from the *D. pulex* transcriptome (Ye *et al*. 2017) in the KEGG Automatic Annotation Server (KAAS) using the GHOSTX program (Moriya *et al*. 2007). A set of 10 135 genes were assigned a KO (KEGG ortholog) number, and from these, 6 282 were assigned to a KEGG pathway map. Hypergeometric tests with Holm-Bonferroni corrected p-values (p-value < 0.05) were used to identify enriched pathways (script available at: https://github.com/Marelize007/RNAseq_obligate_parthenogenesis). The up-and down-regulated genes were analyzed separately to increase the power to detect biologically relevant pathways (Hong *et al*. 2014). The Gene Annotation Easy Viewer (GAEV) (Huynh and Xu 2019) was used to visualize the functional pathways that each gene is mapped to.

### Alternative splicing analysis

Differentially spliced events (DS) were identified using the software rMATS v4.1.1 (Shen *et al*. 2014). Reads mapped to both exons and splice junctions were used to detect the following alternatively spliced events: SE (skipped exon), A5SS (alternative 5’ splice site), A3SS (alternative 3’ splice site), RI (retained intron), and MXE (mutually exclusive exons). An SE event occurs when an entire exon including its flanking introns is spliced out. A5SS and A3SS events may result in the inclusion or exclusion of different parts of exons, while entire introns are retained during RI events. During MXE events only one out of two exons is spliced into the resulting mRNA (Pohl *et al*. 2013, Wang *et al*. 2015). Genes were considered to be differentially spliced if at least four uniquely mapped reads supported the events, reads had a minimum anchor length of 10 nucleotides, the Benjamini-Hochberg adjusted p-value < 0.05, and the difference in exon inclusion level (Δ|Ψ|) > 5% (Shen *et al*. 2014, Suresh *et al*. 2020). Similar to the differential expression analysis, this analysis was completed for each isolate to identify within-isolate differences, as well as by pooling the isolates together to obtain a comprehensive overview of the splicing differences between the early subitaneous and early resting egg production stages. Chi-squared tests were performed using R in Rstudio to test for significant (p < 0.05) over-and-under representation of splicing events within each isolate and the pooled sample.

## Results

### Data quality

Three biological replicates were collected for each isolate during early subitaneous egg and early resting egg production, leading to a total of 18 RNA-seq samples. An average of 26.4 million (SD=5 million) raw reads were sequenced per sample. Our quality check using FastQC revealed no issues related to read quality or adapter contamination. On average 93% (SD = 2%) of the reads uniquely mapped to the *D. pulex* reference genome were retained for differential gene expression and alternative splicing analysis **(Supplementary Table S2)**.

### Differential expression analysis

Mapped read counts were normalized using the regularized log (rlog) transformation function in DEseq2, and a principal component analysis (PCA) was performed to visualize the grouping of samples. The first two principal components accounted for 47% and 17% of the total variance, respectively. The first principal component (PC1) is likely due to differences between the early subitaneous and early resting egg production stages, whereas the second principal component (PC2) could be due to clonal variance **(Figure 3A)**. The same separation pattern for PC1 and PC2 was seen among individual isolates **(Supplementary Figure S1)**. These results strongly suggest that our data captured the transcriptional differences between the early subitaneous and early resting egg production stages and some inter-clonal differences.

**Figure 3.**
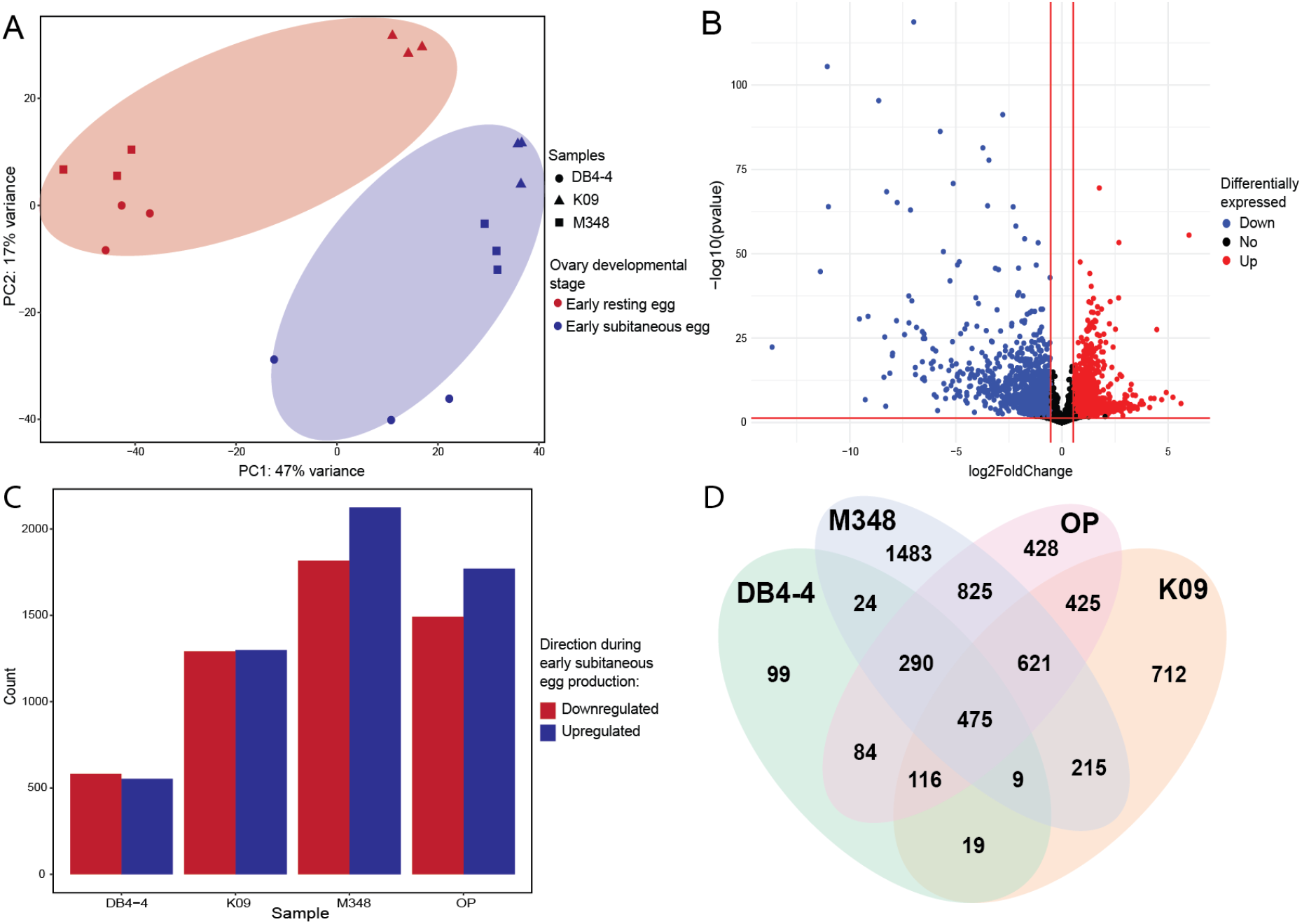
(A) PCA plot based on all samples. (B) Volcano plot of genes differentially expressed during early subitaneous egg production for the pooled sample. (C) Number of genes upregulated and downregulated during early subitaneous egg production in each sample. (D) Venn diagram showing the number of shared differentially expressed genes between early subitaneous and early resting egg production. OP represents the set of differentially expressed genes from the pooled transcriptomic analysis.

To increase statistical power and obtain a comprehensive overview of the main differences between early subitaneous and early resting egg production contributing to PC1, we performed a pooled analysis comparing the developmental stages across all three isolates. A total of 3263 genes were significantly differentially expressed (p-value < 0.05), with 1771 genes upregulated and 1492 genes downregulated in early subitaneous egg production compared to early resting egg production **(Figures 3B and 3C, Supplementary Table S3)**.

To reveal lineage-specific differences and commonalities, the transcriptomes between the two reproductive stages were compared within each isolate. For DB4-4, K09, and M348, a total of 1116, 2591, and 3942 genes were differentially expressed between the two stages, respectively **(Supplementary Table S3)**. Of these genes, 534, 1299 and 2125 were upregulated and 582, 1292, 1817 were downregulated during early subitaneous egg production for DB4-4, K09 and M348, respectively **(Figure 3C)**. Additionally, 475 genes were shared among these isolates and the pooled analysis, with a common set of168 genes upregulated and 307 genes downregulated for each sample. Furthermore, 99, 1483, 712, and 428 genes were uniquely differentially expressed in DB4-4, M348, K09, and the pooled analysis, respectively **(Figure 3D)**.

### KEGG pathway enrichment

A KEGG pathway enrichment analysis comparing the two reproductive stages within each isolate and in the pooled data revealed that the onset of early subitaneous egg production compared to early resting egg production is associated with the upregulation of meiosis and cell cycle genes, as well as the upregulation of genes mapped to sugar, lipid, and hormone metabolic pathways (the same set of genes were downregulated in early resting egg production). On the other hand, downregulated genes in early subitaneous egg production (i.e., upregulated genes in early resting egg production) were mainly enriched in various metabolic, biosynthesis, and signaling pathways **(Figure 4A)**. Specifically, of the genes upregulated during early subitaneous egg production obtained from the pooled analysis, 576 were mapped to KEGG pathways and were enriched in 28 pathways (p.adjust < 0.05, **Supplementary Table S4**). The most notable of the pathways enriched with upregulated genes included the Hedgehog signaling pathway, which plays a vital role in embryonic development by coordinating cell proliferation, coordination, and migration (Carballo *et al*. 2018). Additionally, the cell cycle, oocyte meiosis, and progesterone-mediated oocyte maturation pathways were all significantly enriched for upregulated genes in early subitaneous egg production (i.e., downregulated genes in resting egg production). Significantly enriched sugar and lipid metabolic pathways included starch and sucrose metabolism, galactose metabolism, and sphingolipid metabolism **(Figure 4A)**. KEGG pathways maps for the cell cycle, oocyte meiosis, and progesterone-mediated oocyte maturation pathways were produced using the software GAEV (Huynh and Xu 2019) and can be viewed in **Supplementary Figures 2, 3, and 4**. The same enrichment pattern for the Hedgehog signaling pathway, cell cycle, oocyte meiosis, progesterone-mediated oocyte maturation and various sugar and lipid metabolic pathways observed in the pooled analysis was also identified within individual isolates **(Figure 4B)**.

**Figure 4.**
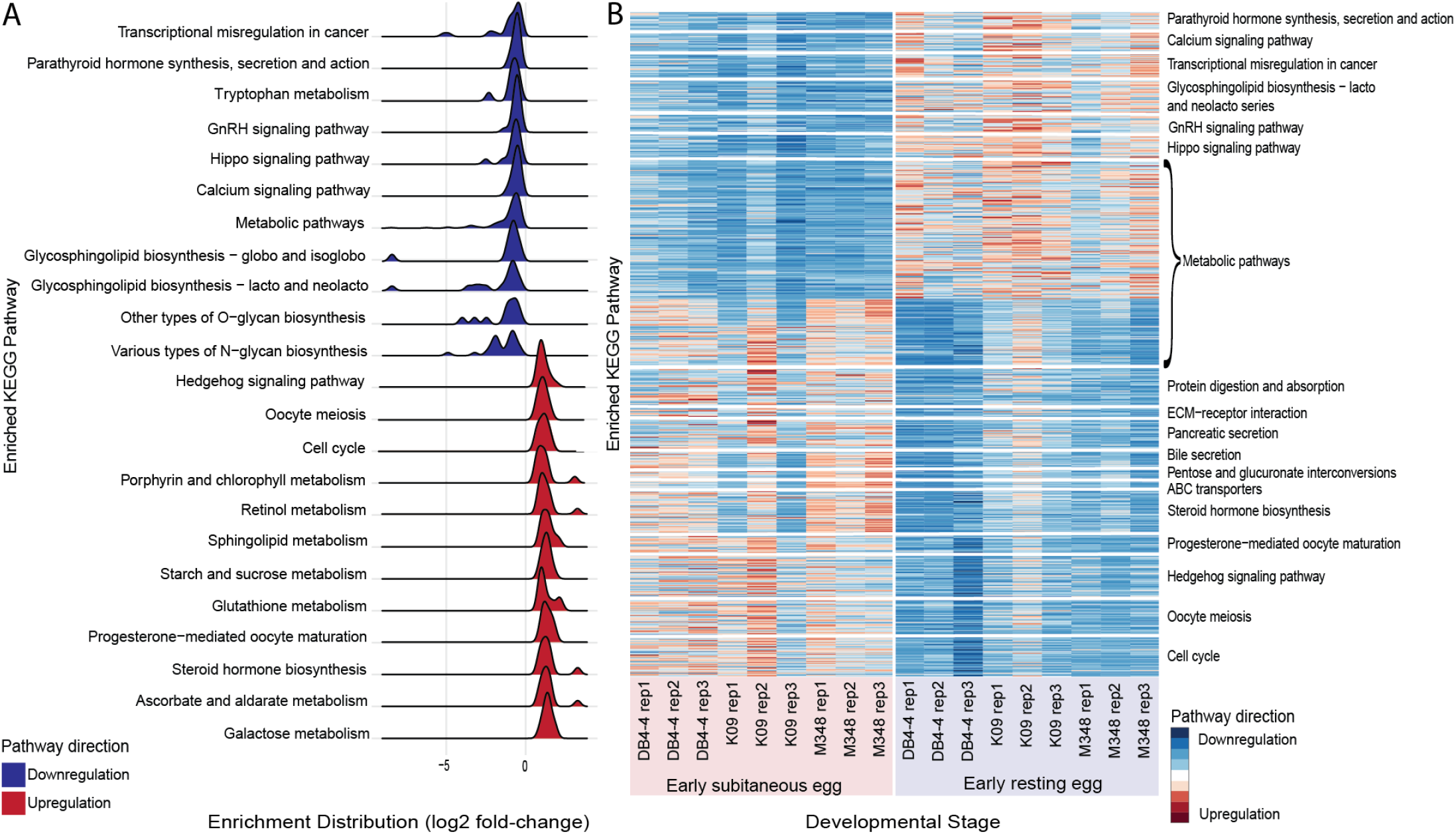
(A) Distribution of up- and downregulated genes during early subitaneous egg production according to log2 fold-change for enriched KEGG pathways. (B) Heatmap illustrating a subset of up- and downregulated KEGG pathways during early subitaneous egg production across all replicates.

Of the downregulated genes for early subitaneous egg production (i.e., upregulated genes in resting egg production) obtained from the pooled analysis, 449 were mapped to KEGG pathways and were enriched in 12 pathways (p.adjust < 0.05, **Supplementary Table S5**). These pathways included metabolic and biosynthesis pathways such as various N-glycan, O-glycan, and glycosphingolipid biosynthesis, as well as the GnRH, Hippo, and calcium signaling pathways **(Figure 4A)**. The GnRH signaling pathway is a regulator of the reproductive system (Kraus *et al*. 2001), while the Hippo signaling pathway controls organ size and development (Boopathy and Hong 2019).

Collectively, these results suggest that the upregulation of meiosis and cell cycle genes and genes mapped to various sugar and lipid metabolic pathways, are associated with early subitaneous egg production. On the opposite, the downregulation of meiosis and cell cycle genes characterizes early resting egg production. Additionally, early subitaneous egg production is associated with a downregulation of genes mapped to various metabolic and biosynthesis pathways including N-glycan, O-glycan, and glycosphingolipid biosynthesis, as well as the GnRH, Hippo, and calcium signaling pathways, whereas all of these pathways are enriched with upregulated genes in early resting egg production.

### GO term enrichment analysis

Our GO term enrichment analysis further corroborated the idea that early subitaneous egg production is associated with an upregulation of meiosis and cell cycle genes and genes mapped to various sugar and lipid metabolic processes, whereas the downregulation of these genes is true of early resting egg production. Upregulated genes in early subitaneous egg production (i.e., downregulated genes in resting egg production) revealed enrichment for GO terms associated with carbohydrate metabolic process (p-value = 0.00014), oogenesis (p-value = 0.0016), lipid transport (p-value = 0.0022), trehalose metabolic process (p-value = 0.0058), regulation of mitotic cell cycle phase transition (p-value = 0.021) and cellular carbohydrate biosynthetic process (p-value = 0.040), **Supplementary Table S6**). In line with the KEGG pathway analysis, our GO term enrichment analysis revealed a downregulation of various metabolic processes in early subitaneous egg production (also meaning upregulation in early resting egg production), including arginine metabolic process (p-value = 0.0065) and proline metabolic process (p-value = 0.0096), cell differentiation (p-value = 0.021), protein glycosylation (p-value = 9 × 10^−7^), and other biosynthetic and signaling processes **(Supplementary Table S7)**. Again, this pattern of regulation observed during early subitaneous egg production was observed within each isolate **(Supplementary Tables S8-S13)**.

### Alternatively spliced genes

Alternatively spliced transcripts were identified between the two stages within each isolate and using a pooled splicing approach to investigate how alternative splicing distinguishes between early subitaneous egg and early resting egg production. Across the pooled samples, 293 differentially spliced transcripts (FDR corrected p-value < 0.05) were identified between early subitaneous egg and early resting egg production. For the individual isolates, 397, 466, and 515 differentially spliced transcripts were identified for DB4-4, M348, and K09, respectively **(Figure 5A, Supplementary Table S14)**. Across all samples analyzed, 60 differentially spliced transcripts were shared **(Figure 5B)**. A GO term enrichment analysis of the differentially spliced transcripts between early subitaneous and early resting egg production shared by at least two samples (total transcripts = 213) showed enrichment for various metabolic and biosynthetic processes including the arginine metabolic process (p-value = 0.043) and prostaglandin metabolic process (p-value = 0.018, **Figure 5C, Supplementary Table S15 – S18**). Additional annotation also revealed the gene UBE2I (UBC9), Ubiquitin Conjugate Enzyme E2 I, to be differentially spliced among all samples. Previous work has shown that UBE2I may fulfil essential roles in regulating gene expression during oocyte growth and maturation (Ihara *et al*. 2008).

**Figure 5.**
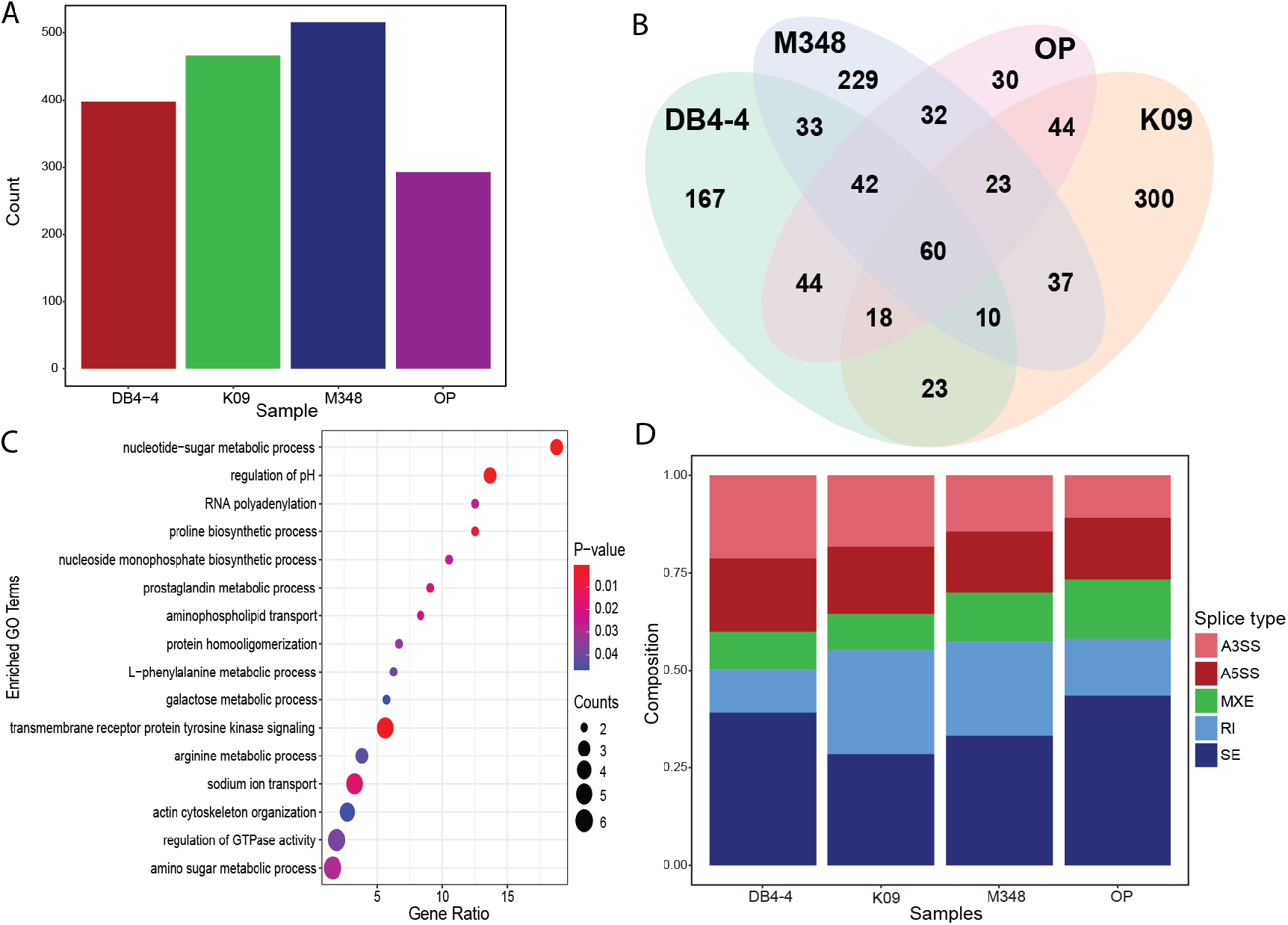
(A) Number of differentially spliced transcripts between early subitaneous and early resting egg production for all samples. (B) Venn diagram showing the number of differentially spliced genes between early subitaneous egg and early resting egg production. OP represents the set of differentially spliced genes from the pooled analysis. (C) Significantly enriched GO terms of differentially spliced genes. (D) Composition of differentially spliced genes.

Furthermore, analyses of the alternative splicing events using single isolates and the pooled samples revealed that skipped exon (SE) events were most abundant, totaling 729 (35%) events across all comparisons, followed by 417 (20%) RI events, 350 A5SS events (17%), 345 A3SS events (17%) and 235 MXE events (11%). **(Figure 5D, Supplementary Table S14)**. Lastly, we tested for over- and under-representation of splicing type among transcripts differentially spliced between early subitaneous and early resting egg production for each isolate and the pooled splicing analysis. The DB4-4 isolate showed an over-representation of A3SS and RI events, while K09 showed an over-representation of RI and SE events (chi-squared test p < 0.05). For the pooled splicing analysis, there was a significant under-representation of A3SS and SE events (chi-squared test p < 0.05), whereas no significant over-representation of splicing events were identified.

### Genes of interest

With early subitaneous egg production showing significant upregulation of meiosis and cell cycle genes (same genes downregulated in early resting egg production), we compiled a list of consensus genes mapped to the following significant pathways: Hedgehog signaling pathway, cell cycle, oocyte meiosis, and progesterone-mediated oocyte maturation pathway. Consensus genes had to be shared by all 3 of the individual isolates or two of the individual isolates and the pooled analysis **(Figure 6)**. These consensus genes mainly consisted of cell cycle regulators such as AURKA, which regulates spindle formation and controls chromosome segregation (Blengini *et al*. 2021), various cyclins including CCNA (cyclin A2), CCNB2 (cyclin B2), CCNB3 (cyclin B3), CLB, and CDC20, CDC4, REC8L, and SMC1. Upregulation across all three isolates and the pooled analysis was observed for paralogs of CDC20 (cell division cycle 20), which activates the anaphase-promotion complex/cyclosome (APC/C) to initiate sister chromatid separation (Jin *et al*. 2010), and paralogs of REC8L, a gene coding the meiosis-specific component of the cohesion complex, which regulates the separation of sister chromatids and homologous chromosomes (Ward *et al*. 2016). Additionally, CDC4 (cell division control protein 4), essential for the transition from G1 to S phase, the onset of anaphase, and the transition from G2 to M phase (Goh and Surana, 1999), showed upregulation in all three isolates and the pooled sample, with DB4-4, M348 and the pooled sample (OP) having an average log2 fold-change > 2.

**Figure 6.**
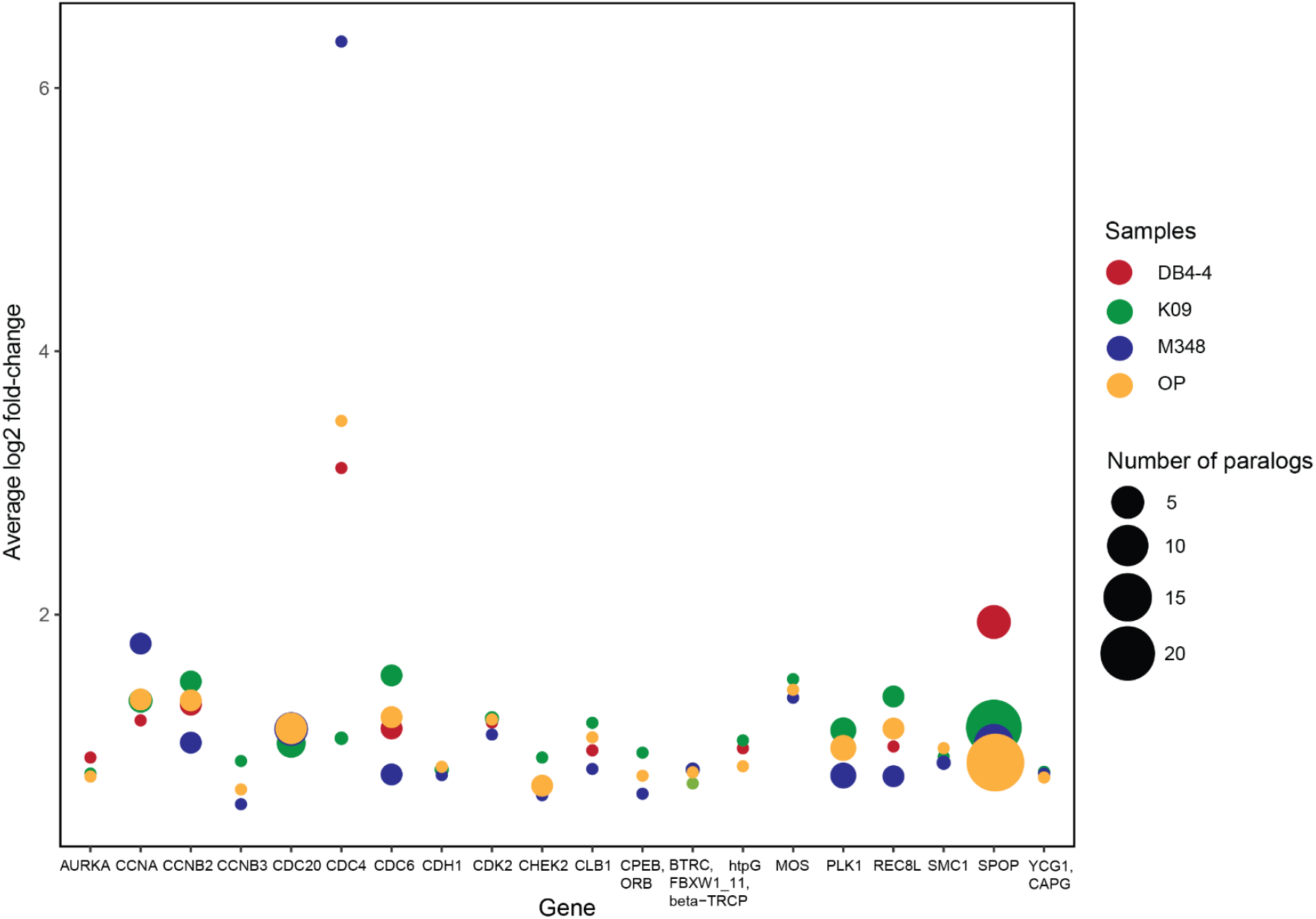
Average log2 fold-change for genes upregulated during early subitaneous embryo production (downregulated in resting egg production) which mapped to the cell cycle, oocyte meiosis, progesterone-mediated oocyte maturation, and Hedgehog signaling pathway. OP represents the pooled sample.

## Discussion

In this study, we investigated the transcriptomic signatures between two modes of parthenogenesis in three obligately parthenogenetic *Daphnia pulex* isolates to gain insight into the genetic modifications critical to the emergence of obligate parthenogenesis. In comparison to previous studies, our results provide a more nuanced view of the origin of obligate parthenogenesis in *Daphnia*.

As stated in the Introduction, previous studies on the transcriptomics of different reproductive stages in *Daphnia* (Xu *et al*. 2022; Huynh *et al*. 2022) suggest that the initiation of parthenogenesis (in subitaneous egg or resting egg production) is associated with an upregulation of metabolic and biosynthesis genes and a downregulation of meiosis and cell cycle genes, leading to the hypothesis that the origin of obligate parthenogenesis in *Daphnia* is due to the extension of parthenogenesis occurring in subitaneous egg production into resting egg production (Xu *et al*. 2022). This hypothesis also draws support from the highly similar cytological observations between the parthenogenesis of subitaneous eggs and resting eggs in *Daphnia* (Zaffagnini and Sabelli 1972).

### Expression of meiosis and cell cycle genes

In the context of this hypothesis, our comparative transcriptomic analysis of the two modes of parthenogenesis (early subitaneous vs. resting egg) in OP *D. pulex* isolates revealed new insights into the genetic modifications associated with obligate parthenogenesis. Our differential gene expression analyses and functional enrichment analyses clearly show that in OP *Daphnia* the parthenogenesis in resting egg production, which distinguishes OP from CP *Daphnia*, is associated with reduced expression of meiosis and cell cycle genes compared to the subitaneous, parthenogenetic egg production. Therefore, despite the highly similar cytological modifications of parthenogenesis in subitaneous egg and resting egg production compared to normal meiosis, the expression requirement for meiosis and cell cycle genes is likely different for the successful execution of these two reproductive stages. Parthenogenesis in resting egg production in OP *Daphnia* may only be achieved with a reduced expression of meiosis and cell cycle genes compared to subitaneous egg production. For the transition from cyclical parthenogenesis to obligate parthenogenesis in *Daphnia* (i.e., meiosis converted to parthenogenesis in resting egg production), substantially reduced expression of genes involved in meiosis and cell cycle seems to be critical, which could be due to introgression of CP *D. pulicaria* alleles into the CP *D. pulex* genomic background (**Figure 7**, Xu *et al*. 2022). We also note that the reduced expression of meiosis and cell cycle genes in resting egg production could be attributed to the fact that a single female *Daphnia* produce a maximum of two resting eggs versus multiple subitaneous eggs.

**Figure 7.**
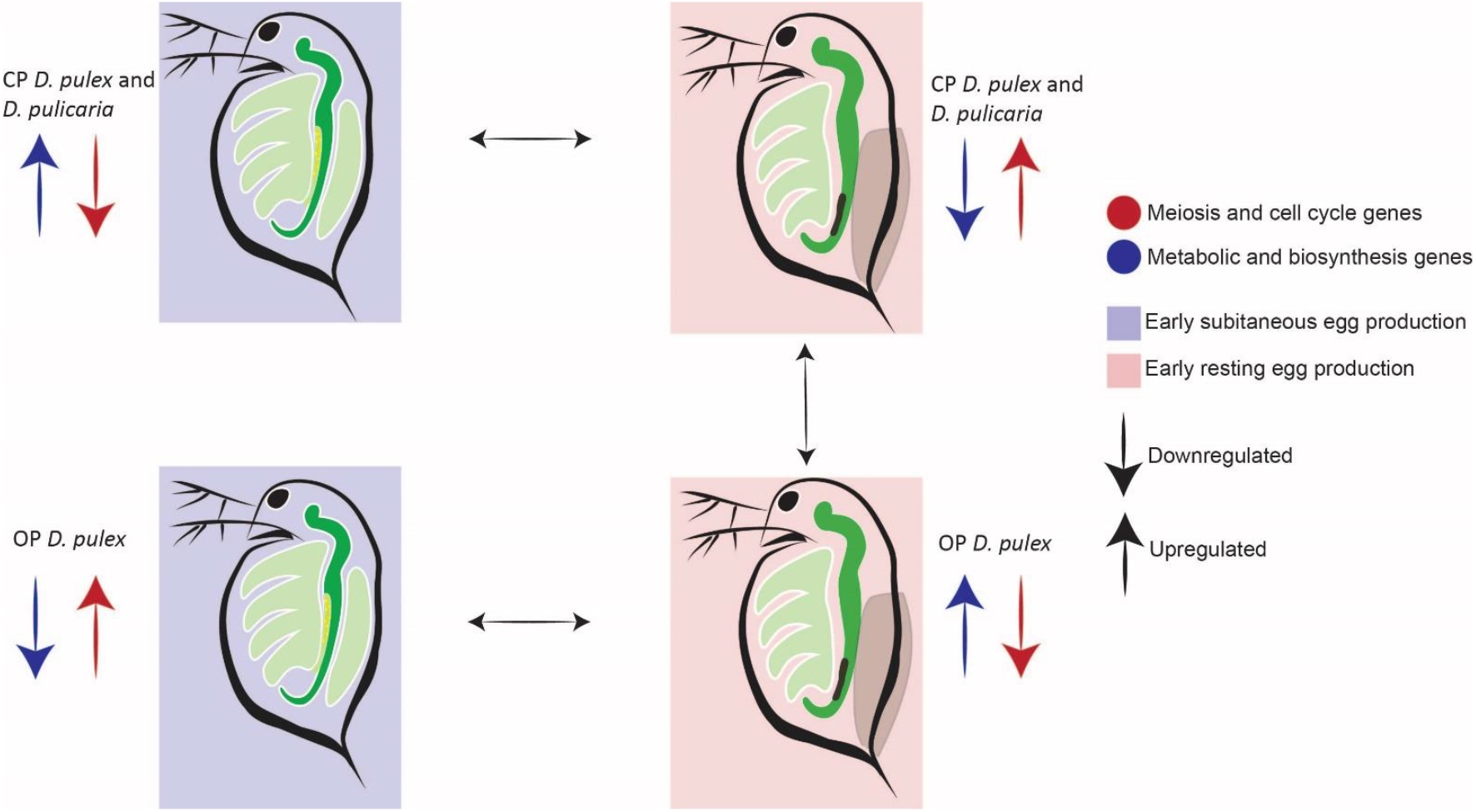
Summary of pathway regulation across various modes of reproduction in OP and CP *Daphnia*.

### Divergent expression of metabolism and biosynthesis genes

Interestingly, our analyses identify a divergent expression pattern of metabolism and biosynthesis pathways in resting egg production vs. subitaneous egg production in OP *Daphnia*. This is worthy of attention because altering the amino acid composition through diet can induce *Daphnia* to enter different reproductive stage (Fink *et al*. 2011, Koch *et al*. 2011), which suggests an important role of metabolism and biosynthesis in regulating the reproduction of *Daphnia*.

In CP *D. pulex* and *D. pulicaria* (Huynh *et al*. 2022), the parental species of OP *Daphnia*, resting egg production is predominantly associated with the downregulation of metabolic and biosynthesis genes. However, the results in this study clearly show that resting egg production in OP *Daphnia* is associated with both the downregulation and upregulation of genes enriched in some specific metabolism and biosynthesis pathways. The identified divergent expression of these pathways may contain the genetic regulators of reproductive modes and indicate the varying metabolic requirements of different reproductive modes in OP *Daphnia*.

### Metabolism and biosynthesis differences between two reproductive modes

The metabolic and biosynthesis genes upregulated during early subitaneous egg production, i.e., downregulated during early resting egg production, were enriched in various sugar metabolic pathways including galactose, starch and sucrose metabolism, sphingolipid metabolism, steroid hormone biosynthesis and progesterone-mediated oocyte maturation **(Figures 4A, B)**. We suggest that a few of these pathways are important for reproductive regulation in crustaceans and arthropods. For example, in the study by Varki *et al*. (2015), it was shown that sphingolipids play a vital role in growth factor signaling and morphogenesis in arthropods, and changes in sphingolipid abundance could impact cell proliferation, apoptosis, senescence, and differentiation (Hannun and Obeid 2002). Moreover, in the brine shrimp, *Artemia franciscana*, that shares reproductive similarities with *CP Daphnia* (Nambu *et al*. 2004), Kojima *et al*. (2013) showed that sphingolipids involved in signaling and signal transduction pathways might be vital in determining which reproductive mode ensue in *A. franciscana* (Kojima *et al*. 2013). Thus, it is likely that sphingolipid metabolism may be critical in initiating and differentiating the two reproductive modes in OP *Daphnia* isolates.

We also note that steroid hormones previously described in crustaceans include ecdysteroids, which play an essential role in regulating the molting cycle, as well as vertebrate-type steroids such as estrogen and progesterone which fulfil functions in vitellogenesis and ovarian development (Lafont and Mathieu 2007, Liu *et al*. 2019, Nakagawa and Henrich 2009). The divergent expression pattern of the steroid hormone biosynthesis pathway and progesterone-mediated oocyte maturation pathway in our study indicates that the two reproductive modes in OP *Daphnia* have differential requirement for the level of steroid hormones. It would be of great interest for future studies to address whether any of the differentially expressed genes in these two pathways drive the initiation of different reproductive modes or their differential expression is a consequence of different reproductive modes.

On the other hand, the metabolic and biosynthesis genes downregulated during early subitaneous egg production, i.e., upregulated during early resting egg production, showed enrichment for various metabolic pathways including arginine metabolism, glycosphingolipid biosynthesis, parathyroid hormone synthesis, secretion, and action **(Figures 4A and B)**. Previous work on the inducibility of resting eggs in CP *Daphnia* isolates have revealed that supplementing specific dietary amino acids such as arginine could promote subitaneous egg production and suppress resting egg production, suggesting their important roles in the switch between parthenogenesis and sexual reproduction in CP *Daphnia* (Fink *et al*. 2011, Koch *et al*. 2011). Intriguingly, out results show the opposite association pattern where the downregulation of genes in the arginine metabolic pathway is associated with early subitaneous egg production and their upregulation is associated with resting egg production **(Supplementary Table S7)**. It is likely that the up- or downregulated expression of the genes in arginine pathway is not equivalent to the increased or reduced availability of arginine to *Daphnia*. On the other hand, it remains to be tested whether the availability of arginine has the different impact on reproductive mode between OP and CP *Daphnia*.

Furthermore, the parathyroid hormone synthesis, secretion and action pathway fulfil functions in regulating the concentrations of calcium, phosphate and active vitamin D metabolites (Murray *et al*. 2005). The upregulation of this pathway indicates another aspect of the special metabolic requirements or consequence for resting egg production in OP *Daphnia*. Notably, because most metabolic pathways were downregulated in early subitaneous egg production and upregulated during early resting egg production, these upregulated metabolic pathways in resting egg production are worthy of special attention in future studies.

### Signaling pathways

We note that resting egg and subitaneous egg production in OP *Daphnia* are also associated with the divergent expression of genes enriched in some important signaling pathways, which may contain some potential genetic triggers of the two distinct reproductive modes. For example, the evolutionarily conserved Hedgehog signaling pathway was enriched with upregulated genes during early subitaneous egg production, and downregulated genes during early resting egg production **(Figures 4A and B)**. This pathway plays an essential role in cell patterning, embryogenesis and development (Carballo *et al*. 2018, Niyaz *et al*. 2019).

Signaling pathways downregulated during early subitaneous egg production and upregulated during early resting egg production provides insight into the transcriptomic signature of resting egg production where OP *Daphnia* shows a major phenotypic modification. These signaling pathways include the calcium signaling pathway, GnRH signaling pathway, and the Hippo signaling pathway **(Figures 4A and B)**. Calcium in *Daphnia* is vital for growth, molting, and ephippia formation (Giardini *et al*. 2015). In contrast to subitaneous egg production, during resting egg formation, *Daphnia* undergo two molting cycles and form an ephippium containing a thick layer of calcium phosphate enclosing the resting eggs (Gerrish and Cáceres 2003). This suggests that the calcium requirement for resting egg production is higher than for subitaneous egg production. Calcium additionally fulfills vital functions in the GnRH signaling pathway through regulating the secretion of the gonadotropins: luteinizing hormone (LH) and follicle stimulating hormone (FSH), both playing a vital role in embryogenesis and reproduction (Haisenleder *et al*. 2003, Marques *et al*. 2000). Lastly, the Hippo signaling pathway which promotes apoptosis and inhibits cell proliferation was also upregulated in early subitaneous egg production and downregulated in early resting egg production (Sherbet 2017).

### Alternative splicing

Our alternative splicing analysis between early subitaneous and early resting egg production revealed 60 differentially spliced transcripts shared by all samples **(Figure 5B)**. Further annotation of the shared differentially spliced transcripts revealed UBE2I (UBC9), Ubiquitin Conjugating Enzyme E2 I, as a gene of interest. Work done in mouse oocytes revealed that disrupted meiotic maturation and defects in spindle organization resulted from the inhibition of UBE2I in fully grown oocytes, while over-expression of UBE2I caused a stimulation of transcription in meiotically incompetent oocytes (Ihara *et al*. 2008, Yuan *et al*. 2014). These results suggest that isoforms of UBE2I, and thus sumoylation, may play an essential role in regulating gene expression during oocyte growth and maturation (Ihara *et al*. 2008).

Additionally, our GO term enrichment analysis of alternatively spliced transcripts shared by at least two samples showed enrichment for various metabolic and biosynthetic processes, including the arginine metabolic process (**Figure 5C**). The enrichment in the arginine metabolic process indicates that alternative splicing may contribute to the regulation of transcript abundance in different reproductive stages in *Daphnia*.

### Candidate genes for obligate parthenogenesis

Lastly, we compiled a list of candidate genes mapped to the cell cycle, oocyte meiosis, progesterone-mediated oocyte maturation, and Hedgehog signaling pathway for future investigation **(Figure 6)**. A gene of particular interest on this list introgressed from *D. pulicaria*, and previously identified as playing an important role in parthenogenesis is CDC20 (Xu *et al*. 2022). During oocyte meiosis, CDC20, a subunit responsible for activating the anaphase-promoting complex/cyclosome (APC/C), promotes progression from metaphase to anaphase via the destruction of cyclin B1 and securin (Jin *et al*. 2010). Studies in mice have shown that a reduction in CDC20 increased the average time from metaphase entry to the onset of anaphase (Jin *et al*. 2010), while a lack of CDC20 caused metaphase arrest (Li *et al*. 2007). This metaphase arrest due to a deficiency in CDC20 was also observed in bovine oocytes (Yang *et al*. 2014) and budding yeast (Lim *et al*. 1998). It will be of great interest to investigate the role of CDC20 in the parthenogenesis of *Daphnia* by manipulating its gene expression (e.g., overexpression).

## Acknowledgements

This work is supported by NIH grant R35GM133730 to SX. We would like to thank the Xu lab members for their helpful discussions.

## Data

The raw reads for this study are deposited at NCBI SRA PRJNA847604.

## Author Contributions

SX designed the study. MS performed the tissue collection, molecular work, and data analysis. SX and MS wrote the manuscript.

