## Supplementary Material for "On the Origin of Obligate Parthenogenesis in *Daphnia pulex*"

**Supplementary Figure S1.** PCA plot depicting the variance between early subitaneous egg and early resting egg production for all three OP *Daphnia pulex* isolates.

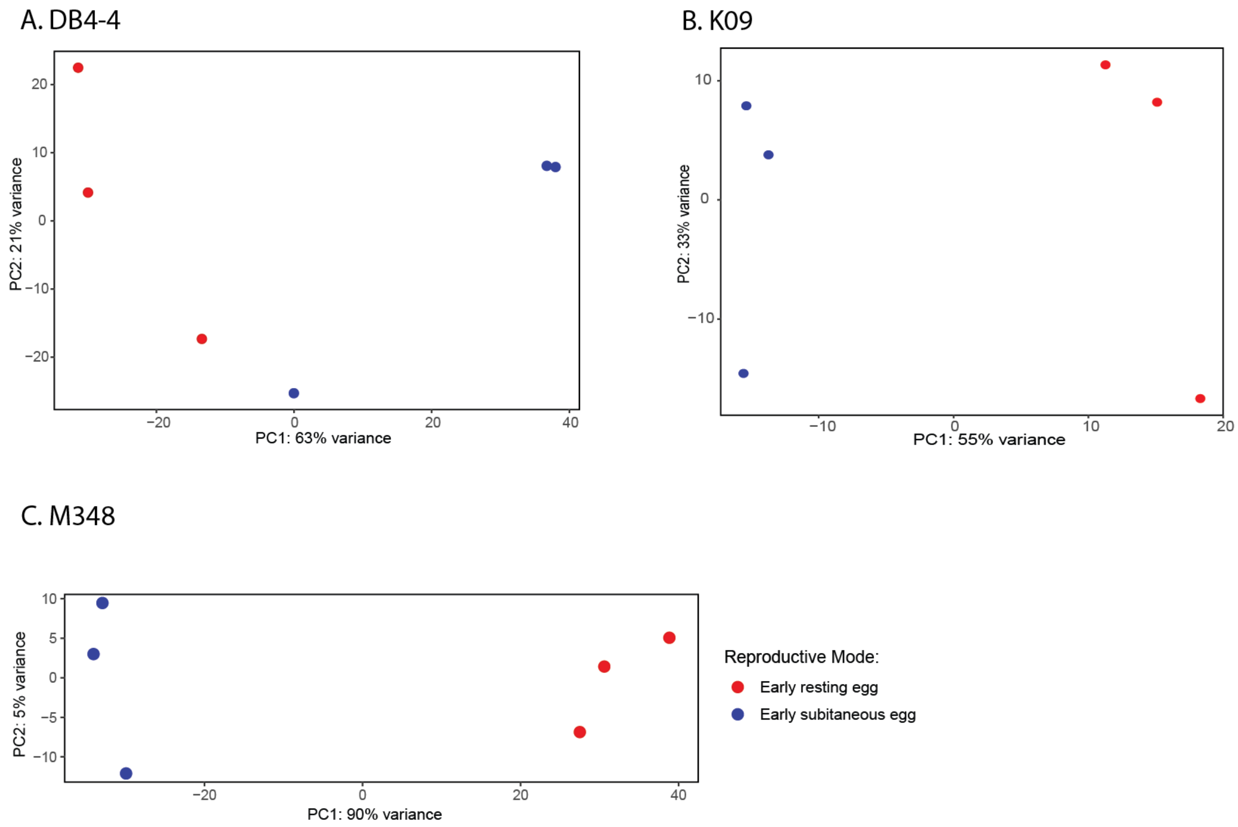

**Supplementary Figure S2.** KEGG pathway enrichment results for the oocyte meiosis pathway using the pooled OP *D. pulex* analysis as input. Genes upregulated in early subitaneous egg production are shown in red, while downregulated genes are shown in blue.

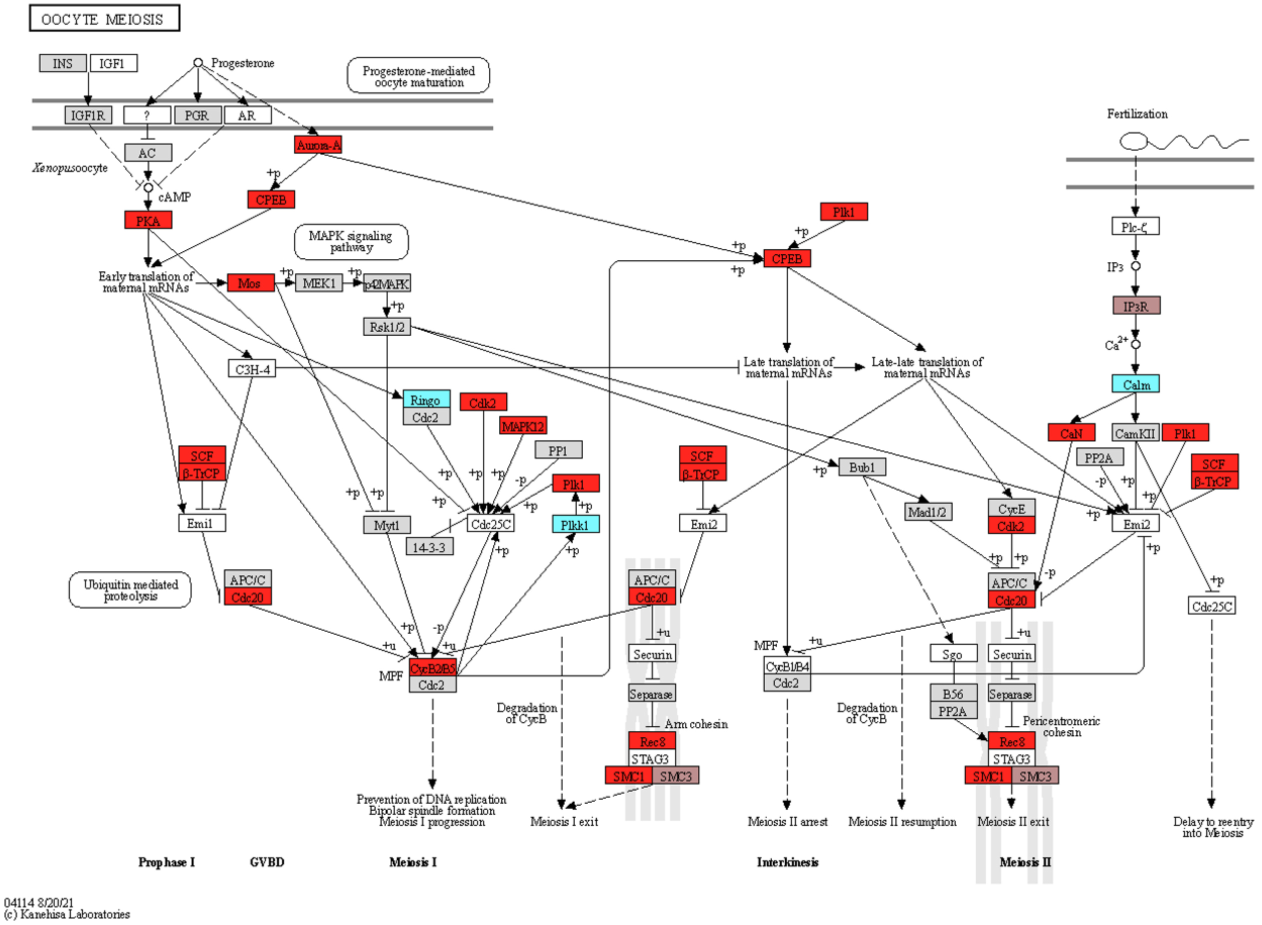

**Supplementary Figure S3.** KEGG pathway enrichment results for the cell-cycle pathway using the pooled OP *D. pulex* analysis as input. Genes upregulated in early subitaneous egg production are shown in red.

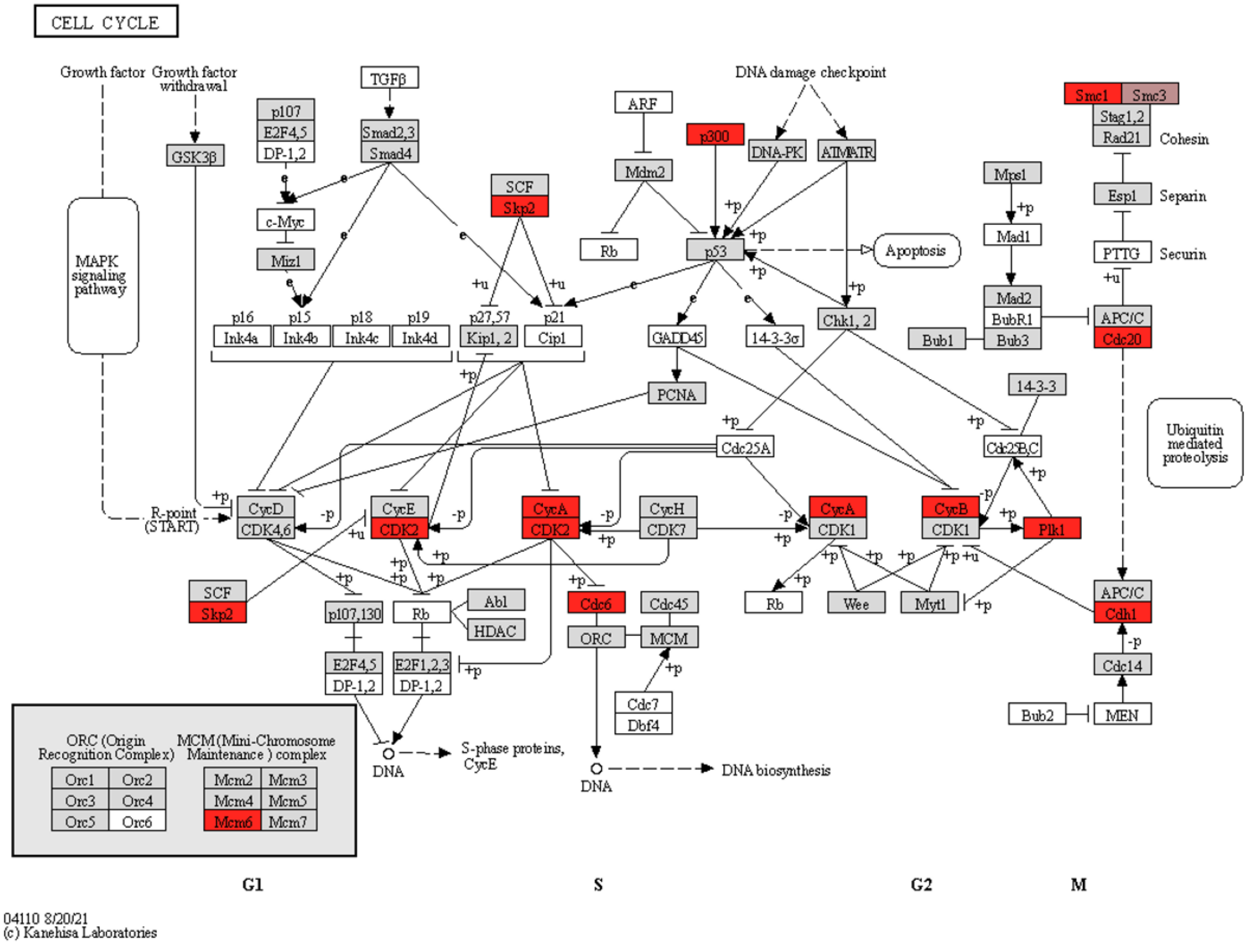

**Supplementary Figure S4.** KEGG pathway enrichment results for the progesterone-mediated oocyte maturation pathway using the pooled OP *D. pulex* analysis as input. Genes upregulated in early subitaneous egg production are shown in red, while downregulated genes are shown in blue.

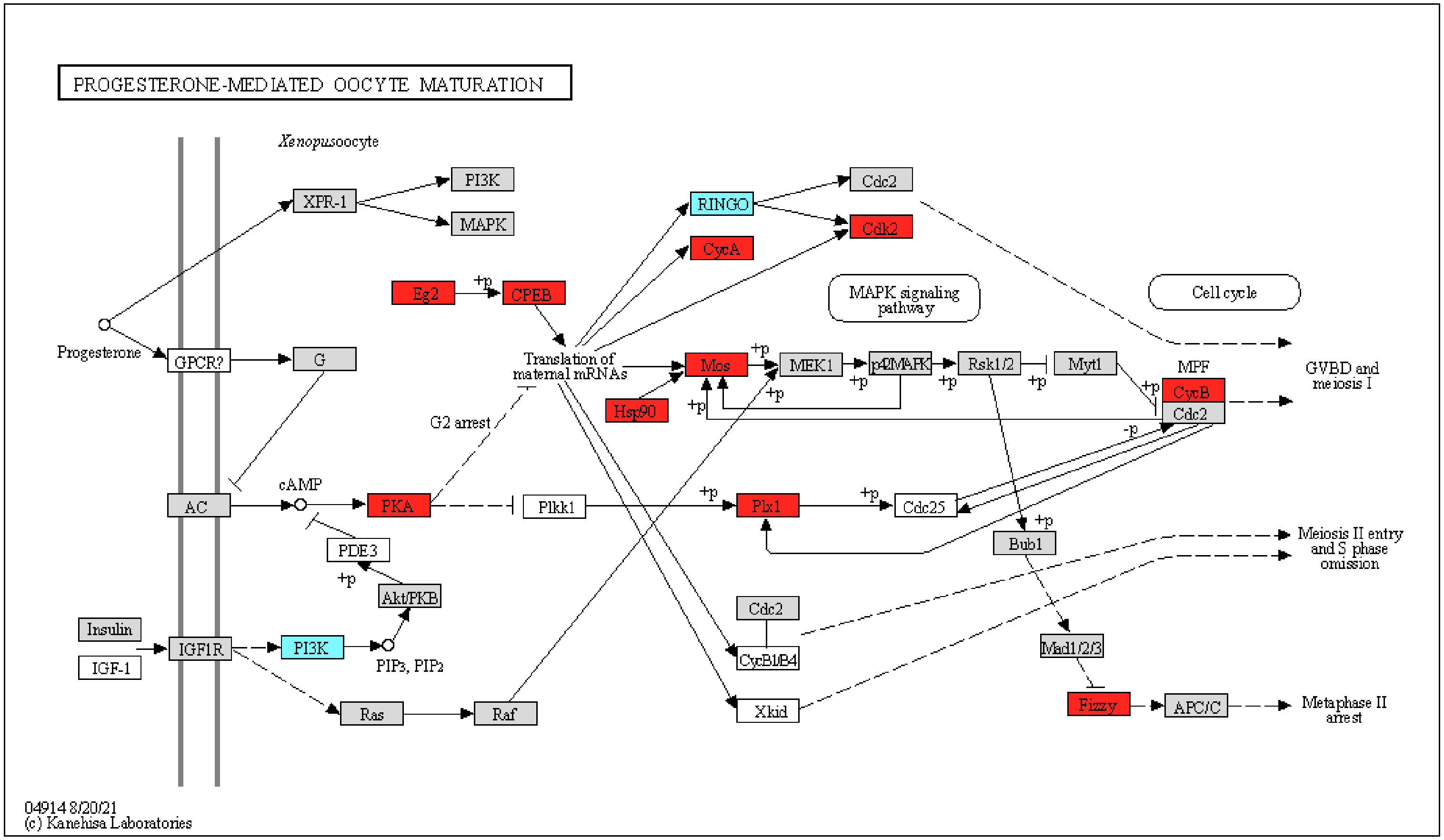

**Supplementary Table S1.** *Daphnia* isolates with collection area.

| Species | Isolates | Location |
|---|---|---|
| *D.pulex (OP)* | K09 | 46°26, -81°3, Kelley Lake, Sudbury, Ontario |
| | DB4-4 | 32° 47’17.6” N, 97° 07’ 27.9” W Drying Bed, Arlington, Texas |
| | Main 348-1 | 42°99, -76°01, Appledore Island, Maine |

**Supplementary Table S2.** Raw read information per replicate and reproductive stage. Samples collected during early subitaneous egg production are labeled as EA, while samples collected during early resting egg production are labeled EM.

| Sample name | Total Sequences | Total Alignments | % Aligned Reads | Assigned Alignments | % Assigned Alignments |
|---|---|---|---|---|---|
| DB4_EA1 | 27845395 | 24815965 | 89.12053501 | 19609955 | 79.00% |
| DB4_EA2 | 29435764 | 27592327 | 93.73742431 | 22212242 | 80.50% |
| DB4_EA3 | 26123157 | 24944712 | 95.4888875 | 20056922 | 80.40% |
| DB4_EM1 | 25187162 | 23723930 | 94.19056422 | 18740622 | 79.00% |
| DB4_EM2 | 23528407 | 20984854 | 89.18943811 | 16423446 | 78.30% |
| DB4_EM3 | 20566563 | 19785077 | 96.20021099 | 15694768 | 79.30% |
| K09_EA1 | 25523596 | 23758018 | 93.08256564 | 18819466 | 79.20% |
| K09_EA2 | 36536128 | 32943316 | 90.16641282 | 25607368 | 77.70% |
| K09_EA3 | 22160525 | 20540182 | 92.68815608 | 16128616 | 78.50% |
| K09_EM1 | 34232440 | 32332770 | 94.4506731 | 24549922 | 75.90% |
| K09_EM2 | 37900528 | 35810972 | 94.48673644 | 27946325 | 78.00% |
| K09_EM3 | 28134572 | 26360685 | 93.69499205 | 20131388 | 76.40% |
| M348_EA1 | 25078184 | 23532340 | 93.83590136 | 18963565 | 80.60% |
| M348_EA2 | 21549151 | 20211120 | 93.79079482 | 16277425 | 80.50% |
| M348_EA3 | 23547136 | 21965139 | 93.28157361 | 17523546 | 79.80% |
| M348_EM1 | 21138421 | 19459647 | 92.05818637 | 14468724 | 74.40% |
| M348_EM2 | 23203998 | 21694488 | 93.49461244 | 16291616 | 75.10% |
| M348_EM3 | 24143875 | 22852688 | 94.65211363 | 17376076 | 76.00% |

**Supplementary Table S3.** Differentially expressed (DE) genes in early subitaneous egg production with direction per isolate and pooled analysis.

| **Isolate** | **Total DE genes** | **Direction** | **Total per Direction** |
|---|---|---|---|
| DB4 | 1115 | Upregulated | 553 |
| | | Downregulated | 582 |
| K09 | 2591 | Upregulated | 1299 |
| | | Downregulated | 1292 |
| M348 | 3942 | Upregulated | 2125 |
| | | Downregulated | 1817 |
| Pooled | 3263 | Upregulated | 1771 |
| | | Downregulated | 1492 |

**Supplementary Table S4**. KEGG pathways significantly upregulated in early subitaneous egg production for the pooled OP *D. pulex* analysis.

| pathway | wht.drawn | wht.in.urn | blk.in.urn | total.draw | p.value | p.adjust |
|---|---|---|---|---|---|---|
| 4341 Hedgehog signaling pathway - fly | 26 | 90 | 6192 | 576 | 6.27904028430267e-08 | 0 |
| 4340 Hedgehog signaling pathway | 25 | 94 | 6188 | 576 | 6.4771192493499e-07 | 1e-04 |
| 0980 Metabolism of xenobiotics by cytochrome P450 | 17 | 54 | 6228 | 576 | 3.37925446287116e-06 | 4e-04 |
| 0982 Drug metabolism - cytochrome P450 | 16 | 51 | 6231 | 576 | 6.907930072752e-06 | 5e-04 |
| 0040 Pentose and glucuronate interconversions | 15 | 46 | 6236 | 576 | 7.86780515332708e-06 | 5e-04 |
| 4974 Protein digestion and absorption | 39 | 207 | 6075 | 576 | 8.46658026184089e-06 | 5e-04 |
| 0500 Starch and sucrose metabolism | 14 | 46 | 6236 | 576 | 3.81206471605243e-05 | 0.0018 |
| 5164 Influenza A | 29 | 146 | 6136 | 576 | 4.50932176972073e-05 | 0.0018 |
| 4976 Bile secretion | 18 | 71 | 6211 | 576 | 4.86636237248086e-05 | 0.0018 |

| | | | | | | |
|---|---|---|---|---|---|---|
| 0053 Ascorbate and aldarate metabolism | 13 | 42 | 6240 | 576 | 5.9223539719106e-05 | 0.002 |
| 0983 Drug metabolism - other enzymes | 18 | 73 | 6209 | 576 | 7.23257925641939e-05 | 0.0022 |
| 0830 Retinol metabolism | 14 | 52 | 6230 | 576 | 0.00016748868216794 | 0.0039 |
| 4512 ECM-receptor interaction | 18 | 78 | 6204 | 576 | 0.00018078553737925 | 0.0039 |
| 5204 Chemical carcinogenesis | 16 | 65 | 6217 | 576 | 0.000184763396136953 | 0.0039 |
| 4114 Oocyte meiosis | 28 | 150 | 6132 | 576 | 0.000188027295251939 | 0.0039 |
| 2010 ABC transporters | 11 | 35 | 6247 | 576 | 0.000188554685724816 | 0.0039 |
| 4110 Cell cycle | 25 | 128 | 6154 | 576 | 0.000198077462046947 | 0.0039 |
| 4972 Pancreatic secretion | 35 | 207 | 6075 | 576 | 0.00024172394619655 | 0.0045 |
| 0860 Porphyrin and chlorophyll metabolism | 12 | 42 | 6240 | 576 | 0.000265753328192558 | 0.0047 |
| 1240 Biosynthesis of cofactors | 28 | 158 | 6124 | 576 | 0.000460124867434089 | 0.0077 |
| 0790 Folate biosynthesis | 10 | 33 | 6249 | 576 | 0.000513144881336937 | 0.0082 |
| 5146 Amoebiasis | 18 | 91 | 6191 | 576 | 0.00129779315335033 | 0.0197 |
| 0052 Galactose metabolism | 12 | 50 | 6232 | 576 | 0.00147655191393492 | 0.0214 |

| | | | | | | |
|---|---|---|---|---|---|---|
| 4914 Progesterone-mediated oocyte maturation | 18 | 95 | 6187 | 576 | 0.0021587295681261 | 0.03 |
| 0480 Glutathione metabolism | 16 | 83 | 6199 | 576 | 0.00311442598430835 | 0.0416 |
| 0140 Steroid hormone biosynthesis | 11 | 48 | 6234 | 576 | 0.00340253101662544 | 0.0421 |
| 0600 Sphingolipid metabolism | 11 | 48 | 6234 | 576 | 0.00340253101662544 | 0.0421 |
| 4111 Cell cycle - yeast | 19 | 108 | 6174 | 576 | 0.0039582896692776 | 0.0472 |

**Supplementary Table S5.** KEGG pathways significantly downregulated in early subitaneous egg production for the pooled OP *D. pulex* analysis.

| pathway | wht.drawn | wht.in.urn | blk.in.urn | total.draw | p.value | p.adjust |
|---|---|---|---|---|---|---|
| 0513 Various types of N-glycan biosynthesis | 25 | 100 | 6182 | 449 | 1.84525706291771e-08 | 0 |
| 1100 Metabolic pathways | 152 | 1491 | 4791 | 449 | 3.01038079929269e-07 | 1e-04 |
| 0601 Glycosphingolipid biosynthesis - lacto and neolacto series | 17 | 69 | 6213 | 449 | 4.52204472130551e-06 | 5e-04 |
| 5202 Transcriptional misregulation in cancer | 27 | 169 | 6113 | 449 | 5.5284548925444e-05 | 0.0047 |
| 0603 Glycosphingolipid biosynthesis - globo and isoglobo series | 11 | 42 | 6240 | 449 | 0.000123527200223239 | 0.0085 |
| 4391 Hippo signaling pathway - fly | 15 | 73 | 6209 | 449 | 0.000157409163045342 | 0.009 |
| 4912 GnRH signaling pathway | 19 | 109 | 6173 | 449 | 0.000222340323313731 | 0.0109 |
| 4020 Calcium signaling pathway | 27 | 192 | 6090 | 449 | 0.000484524928547292 | 0.0208 |
| 4928 Parathyroid hormone synthesis, | 19 | 117 | 6165 | 449 | 0.000564510265219281 | 0.0215 |

| | | | | | | |
|---|---|---|---|---|---|---|
| secretion and action | | | | | | |
| 0514 Other types of O-glycan biosynthesis | 11 | 52 | 6230 | 449 | 0.000908713580803324 | 0.0312 |
| 4392 Hippo signaling pathway - multiple species | 6 | 19 | 6263 | 449 | 0.00156764174079492 | 0.0477 |
| 0380 Tryptophan metabolism | 9 | 40 | 6242 | 449 | 0.00166753562728767 | 0.0477 |

**Supplementary Table S6.** Significantly enriched GO terms upregulated during early subitaneous egg production for the pooled OP *D. pulex* sample.

| **GO.ID** | **Term** | **Annotated** | **Significant** | **Expected** | **weightFisher** | **p.adj** |
|---|---|---|---|---|---|---|
| GO:0006508 | proteolysis | 669 | 80 | 58.17 | 5.2e-05 | 0.070564 |
| GO:0005975 | carbohydrate metabolic process | 275 | 38 | 23.91 | 0.00014 | 0.18984 |
| GO:0055085 | transmembrane transport | 501 | 61 | 43.57 | 0.00129 | 1 |
| GO:0048477 | oogenesis | 16 | 6 | 1.39 | 0.00158 | 1 |
| GO:0006979 | response to oxidative stress | 69 | 14 | 6 | 0.00209 | 1 |
| GO:0006869 | lipid transport | 53 | 11 | 4.61 | 0.00215 | 1 |
| GO:0005991 | trehalose metabolic process | 5 | 3 | 0.43 | 0.00572 | 1 |
| GO:0051493 | regulation of cytoskeleton organization | 26 | 3 | 2.26 | 0.00758 | 1 |
| GO:0071840 | cellular component organization or bioge... | 725 | 46 | 63.04 | 0.0081 | 1 |
| GO:0006606 | protein import into nucleus | 16 | 5 | 1.39 | 0.0095 | 1 |
| GO:0015858 | nucleoside transport | 6 | 3 | 0.52 | 0.01071 | 1 |
| GO:0006541 | glutamine metabolic process | 7 | 3 | 0.61 | 0.01755 | 1 |
| GO:0042554 | superoxide anion generation | 7 | 3 | 0.61 | 0.01755 | 1 |

| | | | | | | |
|---|---|---|---|---|---|---|
| GO:0016998 | cell wall macromolecule catabolic proces... | 32 | 7 | 2.78 | 0.01788 | 1 |
| GO:1901990 | regulation of mitotic cell cycle phase t... | 14 | 5 | 1.22 | 0.0212 | 1 |
| GO:0006030 | chitin metabolic process | 128 | 21 | 11.13 | 0.02214 | 1 |
| GO:0006810 | transport | 1322 | 139 | 114.96 | 0.02326 | 1 |
| GO:0003333 | amino acid transmembrane transport | 20 | 5 | 1.74 | 0.02524 | 1 |
| GO:0006749 | glutathione metabolic process | 27 | 6 | 2.35 | 0.02569 | 1 |
| GO:0051258 | protein polymerization | 36 | 5 | 3.13 | 0.02813 | 1 |
| GO:0006812 | cation transport | 288 | 28 | 25.04 | 0.03532 | 1 |
| GO:0006032 | chitin catabolic process | 29 | 6 | 2.52 | 0.03558 | 1 |
| GO:0034637 | cellular carbohydrate biosynthetic proce... | 9 | 3 | 0.78 | 0.04016 | 1 |
| GO:0007017 | microtubule-based process | 153 | 14 | 13.3 | 0.04478 | 1 |
| GO:0023052 | signaling | 974 | 52 | 84.7 | 0.04552 | 1 |

**Supplementary Table S7.** Significantly enriched GO terms downregulated during early subitaneous egg production for the pooled OP *D. pulex* analysis.

| GO.ID | Term | Annotated | Significant | Expected | weightFisher | p.adj |
|---|---|---|---|---|---|---|
| GO:0006486 | protein glycosylation | 138 | 30 | 10.98 | 9e-07 | 0.0012213 |
| GO:0007156 | homophilic cell adhesion via plasma memb... | 30 | 12 | 2.39 | 1.3e-06 | 0.0017628 |
| GO:0030198 | extracellular matrix organization | 24 | 10 | 1.91 | 6.6e-06 | 0.008943 |
| GO:0007155 | cell adhesion | 114 | 29 | 9.07 | 8.2e-06 | 0.0111028 |
| GO:0006569 | tryptophan catabolic process | 5 | 4 | 0.4 | 0.00019 | 0.25707 |
| GO:0006508 | proteolysis | 669 | 70 | 53.23 | 3e-04 | 0.4056 |
| GO:0018401 | peptidyl-proline hydroxylation to 4-hydr... | 11 | 5 | 0.88 | 0.00096 | 1 |
| GO:0006032 | chitin catabolic process | 29 | 8 | 2.31 | 0.00146 | 1 |
| GO:0006693 | prostaglandin metabolic process | 8 | 4 | 0.64 | 0.00214 | 1 |
| GO:0006355 | regulation of transcription, DNA-templat... | 500 | 53 | 39.78 | 0.00255 | 1 |
| GO:0016998 | cell wall macromolecule catabolic proces... | 32 | 8 | 2.55 | 0.00289 | 1 |

| | | | | | | |
|---|---|---|---|---|---|---|
| GO:0007169 | transmembrane receptor protein tyrosine ... | 33 | 8 | 2.63 | 0.00356 | 1 |
| GO:0019530 | taurine metabolic process | 9 | 4 | 0.72 | 0.00361 | 1 |
| GO:0006691 | leukotriene metabolic process | 9 | 4 | 0.72 | 0.00361 | 1 |
| GO:0006525 | arginine metabolic process | 29 | 7 | 2.31 | 0.00646 | 1 |
| GO:0060429 | epithelium development | 11 | 4 | 0.88 | 0.00832 | 1 |
| GO:0006560 | proline metabolic process | 30 | 7 | 2.39 | 0.00957 | 1 |
| GO:0009435 | NAD biosynthetic process | 7 | 3 | 0.56 | 0.01375 | 1 |
| GO:0007186 | G protein-coupled receptor signaling pat... | 263 | 34 | 20.92 | 0.01664 | 1 |
| GO:0038032 | termination of G protein-coupled recepto... | 8 | 3 | 0.64 | 0.02072 | 1 |
| GO:0030154 | cell differentiation | 51 | 6 | 4.06 | 0.02096 | 1 |
| GO:0006030 | chitin metabolic process | 128 | 22 | 10.18 | 0.02227 | 1 |
| GO:0006555 | methionine metabolic process | 15 | 4 | 1.19 | 0.02667 | 1 |

| | | | | | | |
|---|---|---|---|---|---|---|
| GO:0006979 | response to oxidative stress | 69 | 10 | 5.49 | 0.04501 | 1 |

**Supplementary Table S8.** Significantly enriched GO terms upregulated during early subitaneous egg production for the DB4-4 isolate.

| GO.ID | Term | Annotated | Significant | Expected | weightFisher | p.adj |
|---|---|---|---|---|---|---|
| GO:0051493 | regulation of cytoskeleton organization | 26 | 4 | 0.68 | 0.00067 | 0.90048 |
| GO:0006979 | response to oxidative stress | 69 | 7 | 1.81 | 0.00206 | 1 |
| GO:0051258 | protein polymerization | 36 | 5 | 0.94 | 0.00508 | 1 |
| GO:0005991 | trehalose metabolic process | 5 | 2 | 0.13 | 0.00648 | 1 |
| GO:0006810 | transport | 1322 | 45 | 34.63 | 0.00689 | 1 |
| GO:0048477 | oogenesis | 16 | 3 | 0.42 | 0.0077 | 1 |
| GO:0055085 | transmembrane transport | 501 | 21 | 13.13 | 0.01242 | 1 |
| GO:0003333 | amino acid transmembrane transport | 20 | 3 | 0.52 | 0.01452 | 1 |
| GO:0051301 | cell division | 21 | 3 | 0.55 | 0.01662 | 1 |
| GO:0031667 | response to nutrient levels | 8 | 2 | 0.21 | 0.01723 | 1 |
| GO:0033993 | response to lipid | 39 | 4 | 1.02 | 0.0259 | 1 |
| GO:0009725 | response to hormone | 41 | 4 | 1.07 | 0.02591 | 1 |
| GO:0006955 | immune response | 23 | 3 | 0.6 | 0.02656 | 1 |
| GO:0006508 | proteolysis | 669 | 25 | 17.53 | 0.03 | 1 |
| GO:0006270 | DNA replication initiation | 11 | 2 | 0.29 | 0.03214 | 1 |
| GO:0006749 | glutathione metabolic process | 27 | 3 | 0.71 | 0.03261 | 1 |
| GO:0005975 | carbohydrate metabolic process | 275 | 14 | 7.2 | 0.03276 | 1 |
| GO:0006801 | superoxide metabolic process | 19 | 3 | 0.5 | 0.03759 | 1 |
| GO:0006030 | chitin metabolic process | 128 | 7 | 3.35 | 0.04472 | 1 |

**Supplementary Table S9.** Significantly enriched GO terms downregulated during early subitaneous egg production for the DB4-2 isolate.

| GO.ID | Term | Annotated | Significant | Expected | weightFisher | p.adj |
|---|---|---|---|---|---|---|
| GO:0006486 | protein glycosylation | 138 | 19 | 4.37 | 5.9e-07 | 0.00079296 |
| GO:0007156 | homophilic cell adhesion via plasma memb... | 30 | 6 | 0.95 | 3e-04 | 0.4029 |
| GO:0006693 | prostaglandin metabolic process | 8 | 3 | 0.25 | 0.0016 | 1 |
| GO:0019530 | taurine metabolic process | 9 | 3 | 0.29 | 0.0023 | 1 |
| GO:0006691 | leukotriene metabolic process | 9 | 3 | 0.29 | 0.0023 | 1 |
| GO:0009395 | phospholipid catabolic process | 33 | 5 | 1.05 | 0.0035 | 1 |
| GO:0006979 | response to oxidative stress | 69 | 7 | 2.19 | 0.0059 | 1 |
| GO:0006629 | lipid metabolic process | 260 | 16 | 8.24 | 0.0069 | 1 |
| GO:0007275 | multicellular organism development | 106 | 6 | 3.36 | 0.0123 | 1 |
| GO:0006493 | protein O-linked glycosylation | 7 | 2 | 0.22 | 0.0189 | 1 |
| GO:0038032 | termination of G protein-coupled recepto... | 8 | 2 | 0.25 | 0.0247 | 1 |
| GO:0006508 | proteolysis | 669 | 30 | 21.19 | 0.031 | 1 |
| GO:0006030 | chitin metabolic process | 128 | 9 | 4.05 | 0.0362 | 1 |
| GO:0046339 | diacylglycerol metabolic process | 10 | 2 | 0.32 | 0.038 | 1 |
| GO:0030198 | extracellular matrix organization | 24 | 3 | 0.76 | 0.0389 | 1 |
| GO:0006807 | nitrogen compound metabolic process | 3369 | 112 | 106.71 | 0.0392 | 1 |
| GO:0018401 | peptidyl-proline hydroxylation to 4-hydr... | 11 | 2 | 0.35 | 0.0455 | 1 |

| | | | | | | |
|---|---|---|---|---|---|---|
| GO:0060429 | epithelium development | 11 | 2 | 0.35 | 0.0455 | 1 |

**Supplementary Table S10.** Significantly enriched GO terms upregulated during early subitaneous egg production for the K09 isolate.

| GO.ID | Term | Annotated | Significant | Expected | weightFisher | p.adj |
|---|---|---|---|---|---|---|
| GO:0006270 | DNA replication initiation | 11 | 7 | 0.75 | 1.7e-06 | 0.0022848 |
| GO:0006260 | DNA replication | 76 | 23 | 5.19 | 2.4e-06 | 0.0032232 |
| GO:0006508 | proteolysis | 669 | 72 | 45.65 | 4.5e-06 | 0.006039 |
| GO:0006869 | lipid transport | 53 | 11 | 3.62 | 0.00032 | 0.42912 |
| GO:0006606 | protein import into nucleus | 16 | 6 | 1.09 | 0.00043 | 0.5762 |
| GO:0006541 | glutamine metabolic process | 7 | 4 | 0.48 | 0.00063 | 0.84357 |
| GO:0006338 | chromatin remodeling | 41 | 11 | 2.8 | 0.00169 | 1 |
| GO:0006406 | mRNA export from nucleus | 5 | 3 | 0.34 | 0.00285 | 1 |
| GO:0051493 | regulation of cytoskeleton organization | 26 | 3 | 1.77 | 0.00466 | 1 |
| GO:0071840 | cellular component organization or bioge... | 670 | 44 | 45.72 | 0.00473 | 1 |
| GO:0007093 | mitotic cell cycle checkpoint signaling | 7 | 3 | 0.48 | 0.00898 | 1 |
| GO:1901991 | negative regulation of mitotic cell cycl... | 7 | 3 | 0.48 | 0.00898 | 1 |
| GO:0006334 | nucleosome assembly | 13 | 4 | 0.89 | 0.00932 | 1 |
| GO:0051258 | protein polymerization | 36 | 5 | 2.46 | 0.01241 | 1 |
| GO:1901990 | regulation of mitotic cell cycle phase t... | 14 | 6 | 0.96 | 0.01313 | 1 |
| GO:2000112 | regulation of cellular macromolecule bio... | 556 | 39 | 37.94 | 0.01335 | 1 |
| GO:0030071 | regulation of mitotic metaphase/anaphase... | 8 | 3 | 0.55 | 0.01365 | 1 |
| GO:0051252 | regulation of RNA metabolic process | 553 | 44 | 37.73 | 0.01454 | 1 |
| GO:0048477 | oogenesis | 16 | 4 | 1.09 | 0.02017 | 1 |
| GO:0007017 | microtubule-based process | 153 | 13 | 10.44 | 0.02021 | 1 |

| | | | | | | |
|---|---|---|---|---|---|---|
| GO:0034637 | cellular carbohydrate biosynthetic proce... | 9 | 3 | 0.61 | 0.02535 | 1 |
| GO:0000209 | protein polyubiquitination | 10 | 3 | 0.68 | 0.0264 | 1 |
| GO:0000079 | regulation of cyclin-dependent protein s... | 10 | 3 | 0.68 | 0.0264 | 1 |
| GO:0023052 | signaling | 984 | 47 | 67.14 | 0.02816 | 1 |
| GO:0072488 | ammonium transmembrane transport | 5 | 2 | 0.34 | 0.04047 | 1 |
| GO:0006207 | 'de novo' pyrimidine nucleobase biosynth... | 5 | 2 | 0.34 | 0.04047 | 1 |
| GO:0005991 | trehalose metabolic process | 5 | 2 | 0.34 | 0.04047 | 1 |
| GO:0006979 | response to oxidative stress | 69 | 9 | 4.71 | 0.04364 | 1 |

**Supplementary Table S11.** Significantly enriched GO terms downregulated during early subitaneous egg production for the K09 isolate.

| GO.ID | Term | Annotated | Significant | Expected | weightFisher | p.adj |
|---|---|---|---|---|---|---|
| GO:0006508 | proteolysis | 669 | 85 | 48.22 | 5.5e-09 | 7.392e-06 |
| GO:0007156 | homophilic cell adhesion via plasma memb... | 30 | 13 | 2.16 | 4.7e-08 | 6.3121e-05 |
| GO:0005975 | carbohydrate metabolic process | 275 | 31 | 19.82 | 0.00052 | 0.69784 |
| GO:0006032 | chitin catabolic process | 29 | 8 | 2.09 | 0.00076 | 1 |
| GO:0008152 | metabolic process | 4286 | 325 | 308.95 | 0.00153 | 1 |
| GO:0016998 | cell wall macromolecule catabolic proces... | 32 | 8 | 2.31 | 0.00154 | 1 |
| GO:0009395 | phospholipid catabolic process | 33 | 8 | 2.38 | 0.00191 | 1 |
| GO:0006569 | tryptophan catabolic process | 5 | 3 | 0.36 | 0.00334 | 1 |
| GO:0018401 | peptidyl-proline hydroxylation to 4-hydr... | 11 | 4 | 0.79 | 0.00585 | 1 |
| GO:0006486 | protein glycosylation | 138 | 19 | 9.95 | 0.01219 | 1 |
| GO:0048731 | system development | 62 | 5 | 4.47 | 0.01489 | 1 |
| GO:0048519 | negative regulation of biological proces... | 130 | 5 | 9.37 | 0.01519 | 1 |
| GO:0006633 | fatty acid biosynthetic process | 23 | 5 | 1.66 | 0.02168 | 1 |
| GO:0006979 | response to oxidative stress | 69 | 10 | 4.97 | 0.02506 | 1 |
| GO:0030198 | extracellular matrix organization | 24 | 5 | 1.73 | 0.02581 | 1 |
| GO:0098656 | anion transmembrane transport | 18 | 4 | 1.3 | 0.03042 | 1 |
| GO:0007275 | multicellular organism development | 106 | 12 | 7.64 | 0.03605 | 1 |

| | | | | | | |
|---|---|---|---|---|---|---|
| GO:0072359 | circulatory system development | 5 | 2 | 0.36 | 0.04481 | 1 |
| GO:0006689 | ganglioside catabolic process | 5 | 2 | 0.36 | 0.04481 | 1 |

**Supplementary Table S12.** Significantly enriched GO terms upregulated during early subitaneous egg production for the M348 isolate.

| GO.ID | Term | Annotated | Significant | Expected | weightFisher | p.adj |
|---|---|---|---|---|---|---|
| GO:0005975 | carbohydrate metabolic process | 275 | 70 | 33.87 | 4e-11 | 5.376e-08 |
| GO:0006810 | transport | 1322 | 194 | 162.8 | 5.6e-08 | 7.5208e-05 |
| GO:0055085 | transmembrane transport | 501 | 86 | 61.7 | 3.2e-07 | 0.00042944 |
| GO:0006508 | proteolysis | 669 | 113 | 82.39 | 6.9e-07 | 0.00092529 |
| GO:0006979 | response to oxidative stress | 69 | 24 | 8.5 | 1e-06 | 0.00134 |
| GO:0006635 | fatty acid beta-oxidation | 5 | 5 | 0.62 | 2.8e-05 | 0.037492 |
| GO:0006869 | lipid transport | 53 | 17 | 6.53 | 5.7e-05 | 0.076266 |
| GO:0006030 | chitin metabolic process | 128 | 33 | 15.76 | 1e-04 | 0.1337 |
| GO:0006633 | fatty acid biosynthetic process | 23 | 9 | 2.83 | 0.001 | 1 |
| GO:0006637 | acyl-CoA metabolic process | 13 | 5 | 1.6 | 0.001 | 1 |
| GO:0030206 | chondroitin sulfate biosynthetic process | 5 | 4 | 0.62 | 0.001 | 1 |
| GO:0008152 | metabolic process | 4286 | 537 | 527.82 | 0.0021 | 1 |
| GO:0006040 | amino sugar metabolic process | 140 | 40 | 17.24 | 0.0037 | 1 |
| GO:0006629 | lipid metabolic process | 260 | 56 | 32.02 | 0.0046 | 1 |
| GO:0006027 | glycosaminoglycan catabolic process | 11 | 5 | 1.35 | 0.0068 | 1 |
| GO:0001575 | globoside metabolic process | 8 | 4 | 0.99 | 0.0106 | 1 |
| GO:0016998 | cell wall macromolecule catabolic proces... | 32 | 9 | 3.94 | 0.0126 | 1 |
| GO:1901071 | glucosamine-containing compound metaboli... | 130 | 35 | 16.01 | 0.0145 | 1 |
| GO:0044264 | cellular polysaccharide metabolic proces... | 20 | 5 | 2.46 | 0.0151 | 1 |
| GO:0005991 | trehalose metabolic process | 5 | 3 | 0.62 | 0.0154 | 1 |

| | | | | | | |
|---|---|---|---|---|---|---|
| GO:0071840 | cellular component organization or bioge... | 670 | 39 | 82.51 | 0.017 | 1 |
| GO:0007015 | actin filament organization | 34 | 5 | 4.19 | 0.0262 | 1 |
| GO:0042886 | amide transport | 6 | 3 | 0.74 | 0.0279 | 1 |
| GO:0015858 | nucleoside transport | 6 | 3 | 0.74 | 0.0279 | 1 |
| GO:0018401 | peptidyl-proline hydroxylation to 4-hydr... | 11 | 4 | 1.35 | 0.037 | 1 |
| GO:0048477 | oogenesis | 16 | 5 | 1.97 | 0.0382 | 1 |
| GO:0006812 | cation transport | 289 | 35 | 35.59 | 0.0414 | 1 |
| GO:0009395 | phospholipid catabolic process | 33 | 8 | 4.06 | 0.0428 | 1 |
| GO:0006687 | glycosphingolipid metabolic process | 30 | 11 | 3.69 | 0.0479 | 1 |

**Supplementary Table S13.** Significantly enriched GO terms downregulated during early subitaneous egg production for the M348 isolate.

| GO.ID | Term | Annotated | Significant | Expected | weightFisher | p.adj |
|---|---|---|---|---|---|---|
| GO:0042254 | ribosome biogenesis | 223 | 73 | 22.41 | 5.6e-24 | 7.5264e-21 |
| GO:0006412 | translation | 235 | 69 | 23.62 | 9.6e-19 | 1.28928e-15 |
| GO:0006555 | methionine metabolic process | 15 | 8 | 1.51 | 3.4e-05 | 0.045628 |
| GO:0006730 | one-carbon metabolic process | 31 | 11 | 3.12 | 0.00013 | 0.17433 |
| GO:0030198 | extracellular matrix organization | 24 | 9 | 2.41 | 0.00032 | 0.4288 |
| GO:0007156 | homophilic cell adhesion via plasma memb... | 30 | 10 | 3.02 | 0.00046 | 0.61594 |
| GO:0006569 | tryptophan catabolic process | 5 | 4 | 0.5 | 0.00047 | 0.62886 |
| GO:0000097 | sulfur amino acid biosynthetic process | 5 | 4 | 0.5 | 0.00047 | 0.62886 |
| GO:0006486 | protein glycosylation | 138 | 28 | 13.87 | 0.00098 | 1 |
| GO:0006518 | peptide metabolic process | 272 | 78 | 27.34 | 0.00299 | 1 |
| GO:0006807 | nitrogen compound metabolic process | 3369 | 371 | 338.6 | 0.00405 | 1 |
| GO:0007601 | visual perception | 17 | 6 | 1.71 | 0.00473 | 1 |
| GO:0007155 | cell adhesion | 114 | 24 | 11.46 | 0.00479 | 1 |
| GO:0006123 | mitochondrial electron transport, cytoch... | 8 | 4 | 0.8 | 0.00509 | 1 |
| GO:0042157 | lipoprotein metabolic process | 29 | 5 | 2.91 | 0.00867 | 1 |
| GO:0006030 | chitin metabolic process | 128 | 21 | 12.86 | 0.00899 | 1 |
| GO:0009084 | glutamine family amino acid biosynthetic... | 12 | 4 | 1.21 | 0.016 | 1 |
| GO:0018401 | peptidyl-proline hydroxylation to 4-hydr... | 11 | 4 | 1.11 | 0.01876 | 1 |

| | | | | | | |
|---|---|---|---|---|---|---|
| GO:0042537 | benzene-containing compound metabolic pr... | 11 | 4 | 1.11 | 0.01876 | 1 |
| GO:0006525 | arginine metabolic process | 29 | 7 | 2.91 | 0.0219 | 1 |
| GO:0035235 | ionotropic glutamate receptor signaling ... | 17 | 5 | 1.71 | 0.02243 | 1 |
| GO:0006493 | protein O-linked glycosylation | 7 | 3 | 0.7 | 0.02597 | 1 |
| GO:0044093 | positive regulation of molecular functio... | 63 | 6 | 6.33 | 0.02624 | 1 |
| GO:0006560 | proline metabolic process | 30 | 7 | 3.02 | 0.02796 | 1 |
| GO:0006414 | translational elongation | 18 | 4 | 1.81 | 0.0282 | 1 |
| GO:0006805 | xenobiotic metabolic process | 17 | 3 | 1.71 | 0.02827 | 1 |
| GO:0007602 | phototransduction | 18 | 5 | 1.81 | 0.02856 | 1 |
| GO:0035249 | synaptic transmission, glutamatergic | 18 | 5 | 1.81 | 0.02856 | 1 |
| GO:0007218 | neuropeptide signaling pathway | 13 | 4 | 1.31 | 0.03457 | 1 |
| GO:0015671 | oxygen transport | 8 | 3 | 0.8 | 0.0385 | 1 |
| GO:0006693 | prostaglandin metabolic process | 8 | 3 | 0.8 | 0.0385 | 1 |
| GO:0007186 | G protein-coupled receptor signaling pat... | 259 | 36 | 26.03 | 0.03968 | 1 |

**Supplementary Table S14.** The number of differentially spliced events between early subitaneous egg and early resting egg production per isolate and pooled sample.

| **Comparison** | **A3SS** | **A5SS** | **RI** | **SE** | **MXE** | **Total** |
|---|---|---|---|---|---|---|
| DB4-4 | 104 | 92 | 54 | 191 | 47 | 488 |
| M348 | 84 | 90 | 139 | 193 | 74 | 580 |
| K09 | 115 | 108 | 168 | 179 | 56 | 626 |
| Pooled | 42 | 60 | 56 | 166 | 58 | 382 |
| Total (Percentage) | 345 (16.6) | 350 (16.9) | 417 (20.1) | 729 (35.1) | 235 (11.3) | 2076 |

**Supplementary Table S15.** GO terms significantly enriched with alternatively spliced transcripts during early subitaneous egg production for the pooled analysis.

| GO.ID | Term | Annotated | Significant | Expected | weightFisher | p.adj |
|---|---|---|---|---|---|---|
| GO:0009225 | nucleotide-sugar metabolic process | 6 | 3 | 0.16 | 0.00037 | 0.50209 |
| GO:0006885 | regulation of pH | 8 | 3 | 0.22 | 0.00099 | 1 |
| GO:0007169 | transmembrane receptor protein tyrosine kinase signaling | 33 | 5 | 0.89 | 0.00176 | 1 |
| GO:0006561 | proline biosynthetic process | 6 | 2 | 0.16 | 0.01017 | 1 |
| GO:0006814 | sodium ion transport | 56 | 5 | 1.52 | 0.01722 | 1 |
| GO:0006693 | prostaglandin metabolic process | 8 | 2 | 0.22 | 0.01832 | 1 |
| GO:0015917 | aminophospholipid transport | 9 | 2 | 0.24 | 0.02314 | 1 |
| GO:0043631 | RNA polyadenylation | 6 | 2 | 0.16 | 0.02694 | 1 |
| GO:0009124 | nucleoside monophosphate biosynthetic process | 7 | 2 | 0.19 | 0.02694 | 1 |
| GO:0006040 | amino sugar metabolic process | 140 | 6 | 3.79 | 0.02827 | 1 |
| GO:0051260 | protein homooligomerization | 11 | 2 | 0.3 | 0.03412 | 1 |
| GO:0043087 | regulation of GTPase activity | 117 | 6 | 3.17 | 0.03858 | 1 |
| GO:0006558 | L-phenylalanine metabolic process | 12 | 2 | 0.32 | 0.04023 | 1 |
| GO:0006525 | arginine metabolic process | 29 | 3 | 0.78 | 0.04255 | 1 |
| GO:0030036 | actin cytoskeleton organization | 54 | 4 | 1.46 | 0.04614 | 1 |
| GO:0006012 | galactose metabolic process | 13 | 2 | 0.35 | 0.04671 | 1 |

**Supplementary Table S16.** GO terms significantly enriched with alternatively spliced transcripts during early subitaneous egg production for the K09 isolate.

| GO.ID | Term | Annotated | Significant | Expected | weightFisher | p.adj |
|---|---|---|---|---|---|---|
| GO:0009225 | nucleotide-sugar metabolic process | 6 | 3 | 0.21 | 0.00074 | 1 |
| GO:0006885 | regulation of pH | 8 | 3 | 0.27 | 0.00196 | 1 |
| GO:0018401 | peptidyl-proline hydroxylation to 4-hydr... | 11 | 3 | 0.38 | 0.00536 | 1 |
| GO:0034654 | nucleobase-containing compound biosynthe... | 729 | 24 | 25.01 | 0.0069 | 1 |
| GO:0006525 | arginine metabolic process | 29 | 4 | 0.99 | 0.01634 | 1 |
| GO:0006493 | protein O-linked glycosylation | 7 | 2 | 0.24 | 0.02196 | 1 |
| GO:0007169 | transmembrane receptor protein tyrosine ... | 33 | 4 | 1.13 | 0.02532 | 1 |
| GO:0038032 | termination of G protein-coupled recepto... | 8 | 2 | 0.27 | 0.02863 | 1 |
| GO:0003333 | amino acid transmembrane transport | 20 | 3 | 0.69 | 0.02953 | 1 |
| GO:0043631 | RNA polyadenylation | 6 | 2 | 0.21 | 0.03419 | 1 |
| GO:0009124 | nucleoside monophosphate biosynthetic pr... | 7 | 2 | 0.24 | 0.03419 | 1 |
| GO:0006006 | glucose metabolic process | 32 | 2 | 1.1 | 0.03432 | 1 |
| GO:0006814 | sodium ion transport | 56 | 5 | 1.92 | 0.04213 | 1 |
| GO:0006040 | amino sugar metabolic process | 140 | 4 | 4.8 | 0.04485 | 1 |
| GO:0006560 | proline metabolic process | 30 | 4 | 1.03 | 0.04705 | 1 |
| GO:0006821 | chloride transport | 24 | 3 | 0.82 | 0.04745 | 1 |

**Supplementary Table S17.** GO terms significantly enriched with alternatively spliced transcripts during early subitaneous egg production for the M348 isolate.

| GO.ID | Term | Annotated | Significant | Expected | weightFisher | p.adj |
|---|---|---|---|---|---|---|
| GO:0006885 | regulation of pH | 8 | 4 | 0.25 | 6.2e-05 | 0.084134 |
| GO:0006012 | galactose metabolic process | 13 | 4 | 0.41 | 0.00056 | 0.75936 |
| GO:0009225 | nucleotide-sugar metabolic process | 6 | 3 | 0.19 | 0.00058 | 0.7859 |
| GO:0006000 | fructose metabolic process | 14 | 4 | 0.44 | 0.00076 | 1 |
| GO:0006096 | glycolytic process | 26 | 5 | 0.82 | 0.00116 | 1 |
| GO:0006094 | gluconeogenesis | 31 | 5 | 0.98 | 0.00263 | 1 |
| GO:0006013 | mannose metabolic process | 23 | 4 | 0.73 | 0.00538 | 1 |
| GO:0006739 | NADP metabolic process | 20 | 5 | 0.63 | 0.0056 | 1 |
| GO:0007059 | chromosome segregation | 30 | 3 | 0.95 | 0.00573 | 1 |
| GO:1901135 | carbohydrate derivative metabolic proces... | 474 | 26 | 15 | 0.00854 | 1 |
| GO:0046785 | microtubule polymerization | 5 | 2 | 0.16 | 0.00936 | 1 |
| GO:0006796 | phosphate-containing compound metabolic ... | 843 | 34 | 26.68 | 0.01201 | 1 |
| GO:0006098 | pentose-phosphate shunt | 16 | 3 | 0.51 | 0.0129 | 1 |
| GO:0140014 | mitotic nuclear division | 32 | 3 | 1.01 | 0.01373 | 1 |
| GO:0019637 | organophosphate metabolic process | 269 | 15 | 8.51 | 0.01804 | 1 |
| GO:0051656 | establishment of organelle localization | 7 | 2 | 0.22 | 0.01885 | 1 |
| GO:0007169 | transmembrane receptor protein tyrosine ... | 33 | 4 | 1.04 | 0.01945 | 1 |
| GO:0043087 | regulation of GTPase activity | 117 | 7 | 3.7 | 0.02165 | 1 |
| GO:0055085 | transmembrane transport | 501 | 20 | 15.86 | 0.02175 | 1 |
| GO:0006693 | prostaglandin metabolic process | 8 | 2 | 0.25 | 0.02461 | 1 |

| | | | | | | |
|---|---|---|---|---|---|---|
| GO:0046497 | nicotinate nucleotide metabolic process | 8 | 2 | 0.25 | 0.02461 | 1 |
| GO:0034613 | cellular protein localization | 230 | 8 | 7.28 | 0.02484 | 1 |
| GO:0010970 | transport along microtubule | 9 | 2 | 0.28 | 0.03099 | 1 |
| GO:0016477 | cell migration | 9 | 2 | 0.28 | 0.03099 | 1 |
| GO:0015917 | aminophospholipid transport | 9 | 2 | 0.28 | 0.03099 | 1 |
| GO:1902850 | microtubule cytoskeleton organization in... | 9 | 2 | 0.28 | 0.03099 | 1 |
| GO:0009124 | nucleoside monophosphate biosynthetic pr... | 7 | 2 | 0.22 | 0.03153 | 1 |
| GO:1902600 | proton transmembrane transport | 21 | 2 | 0.66 | 0.03159 | 1 |
| GO:0006869 | lipid transport | 53 | 6 | 1.68 | 0.0387 | 1 |

**Supplementary Table S18.** GO terms significantly enriched with alternatively spliced transcripts during early subitaneous egg production for the DB4-4 isolate.

| GO.ID | Term | Annotated | Significant | Expected | weightFisher | p.adj |
|---|---|---|---|---|---|---|
| GO:0006885 | regulation of pH | 8 | 3 | 0.2 | 0.00084 | 1 |
| GO:0044093 | positive regulation of molecular functio... | 64 | 3 | 1.64 | 0.00622 | 1 |
| GO:0006094 | gluconeogenesis | 31 | 4 | 0.79 | 0.00759 | 1 |
| GO:0006561 | proline biosynthetic process | 6 | 2 | 0.15 | 0.00912 | 1 |
| GO:0051090 | regulation of DNA-binding transcription ... | 6 | 2 | 0.15 | 0.00912 | 1 |
| GO:0072666 | establishment of protein localization to... | 6 | 2 | 0.15 | 0.00912 | 1 |
| GO:0006693 | prostaglandin metabolic process | 8 | 2 | 0.2 | 0.01646 | 1 |
| GO:0006090 | pyruvate metabolic process | 35 | 5 | 0.9 | 0.02029 | 1 |
| GO:0016477 | cell migration | 9 | 2 | 0.23 | 0.02082 | 1 |
| GO:0015917 | aminophospholipid transport | 9 | 2 | 0.23 | 0.02082 | 1 |
| GO:0006040 | amino sugar metabolic process | 140 | 5 | 3.58 | 0.02568 | 1 |
| GO:0006096 | glycolytic process | 26 | 3 | 0.67 | 0.02779 | 1 |
| GO:0051260 | protein homooligomerization | 11 | 2 | 0.28 | 0.03076 | 1 |
| GO:0006888 | endoplasmic reticulum to Golgi vesicle-m... | 13 | 2 | 0.33 | 0.04219 | 1 |
| GO:0000910 | cytokinesis | 14 | 2 | 0.36 | 0.04841 | 1 |
